# Genome-wide CRISPR screen reveals PEX11B as a host restriction factor against ORFV through membrane fluidity regulation

**DOI:** 10.1101/2025.11.28.691156

**Authors:** Xiaoran Gao, Jinyang Hao, Sha Lu, Shida Wang, Yu Sun, Xin Ke, Xia Gao, Yuan Su, Yuze Sun, Yu Tian, Wenyu Yan, Jinliang Wang, Rong Hai, Qianyi Zhang, Jingfei Wang, Wei Hu, Guojun Wang

## Abstract

Host-pathogen interactions are shaped by cellular restriction factors that direct antiviral defenses. We built the first ovine genome-wide CRISPR knockout library in sheep testis (OA3.Ts) cells, targeting all protein-coding genes. Using this platform, we identified PEX11B, a peroxisomal membrane regulatory protein, as a strong restriction factor against orf virus (ORFV) infection. Removing PEX11B increased viral susceptibility and triggered severe cytopathic effects with membrane fusion and syncytia formation. Mechanistic studies showed that PEX11B knockout harmed peroxisomal integrity and disrupted lipid metabolism. This led to greater plasma membrane fluidity, creating a proviral environment that allowed more viral entry and replication. These results reveal a new antiviral function for PEX11B in blocking viral infection and underscore the importance of peroxisomal regulation in host-virus interactions.

## 1 Introduction

Orf virus (ORFV), responsible for contagious pustular dermatitis (also known as ecthyma contagiosum), was first discovered in 1920 and is now present in more than 70 countries across Africa, the Near and Middle East, and Asia, leading to significant economic losses each year.[1–4]. Annually, ORFV infects about 2 million small ruminants worldwide, causing an overall death rate of 10–20%, which rises to over 90% in lambs under three months old. The virus often triggers outbreaks even among fully vaccinated herds[1, 5–7]. Since 2017, large-scale epidemic waves have been documented in Uruguay, Colombia, Morocco and Brazil, underscoring an expanding threat to animal husbandry and zoonotic public health [1, 8–11]. The substantial economic burden imposed by ORFV, which encompasses production losses, treatment costs, and trade restrictions, underscores the imperative need for a nuanced understanding of the molecular mechanisms governing its infection, pathogenesis, and interaction with the host immune system [3, 12].

The ORFV infection cycle initiates with viral attachment to as-yet-unidentified host cell receptors, followed by entry and cytoplasmic replication via the formation of distinctive intracytoplasmic viral factories [13, 14]. Following genome replication and virion assembly, progeny virions are released through lytic or non-lytic mechanisms. Intriguingly, ORFV exhibits broad tropism, infecting diverse cell types across multiple species, including sheep, goats, humans, and other mammals, suggesting the exploitation of conserved or multifactorial entry pathways. [7, 11, 15, 16]. Recent research has begun to uncover the viral components involved in ORFV entry and replication. However, the host cell factors and mechanisms that play roles in these processes remain largely unexplored [17, 18]. Unraveling these virus-host interactions could unveil novel targets for antiviral intervention, offering a strategic advantage over traditional approaches that focus solely on viral proteins, as host-directed therapies may reduce the likelihood of resistance emergence.

Genetic screening technologies have revolutionized the identification of host dependencies critical for viral replication, providing actionable insights for antiviral development [19, 20]. Early methodologies, such as RNA interference (RNAi) and insertional mutagenesis, enabled large-scale functional genomics screens [21, 22]. More recently, CRISPR/Cas9-based genome-wide knockout screens have emerged as a powerful tool for systematically interrogating host factors essential for infections caused by SARS-CoV-2, Ebola virus, West Nile virus, and other high-consequence pathogens [23–30]. Notably, analogous CRISPR screens in livestock species such as pigs challenged with Japanese encephalitis virus, cattle with Bovine Parainfluenza Virus Type 3, and poultry with Avian influenza virus have successfully identified host determinants of viral [31–33]. However, despite the agricultural and economic significance of sheep, genome-wide CRISPR screens in ovine models remain conspicuously absent, primarily due to the lack of immortalized, CRISPR-compatible ovine cell lines. Addressing this gap is essential for advancing our understanding of ORFV pathogenesis and host adaptation.

Here, we address this gap by establishing OA3.Ts cells stably expressing Cas9 and performing the first ovine genome-wide CRISPR/Cas9 screen to identify host restriction factors against ORFV. Our screen pinpointed PEX11B, a peroxisomal membrane regulator, as a potent antiviral factor. Genetic ablation of PEX11B dramatically enhanced ORFV susceptibility, inducing cytopathic effects characterized by membrane fusion and syncytia formation. Mechanistically, PEX11B knockout disrupted peroxisomal integrity, dysregulating lipid metabolism and increasing plasma membrane fluidity—a proviral microenvironment facilitating viral entry and replication. These findings unveil PEX11B as a novel restriction factor and underscore peroxisomal regulation as a pivotal determinant of host-virus interactions, offering a platform for targeted interventions in livestock health.

## 2 Results

### 2.1 Development of an immortalized OA3.Ts-Cas9 cell line for genome-wide CRISPR screening

Despite the transformative potential of CRISPR screening in *Ovis* species for studying mammalian physiology and disease, progress has been hindered by the lack of expandable cell lines suitable for large-scale genetic interrogation. To address this bottleneck, we established an immortalized sheep testis (OA3.Ts) cell line stably expressing Cas9 and human telomerase (hTERT).

We engineered a lentiviral construct encoding hTERT—fused with P2A, NLS, and HA tags—and inserted it into a linearized lenti-blast-Cas9 plasmid via BamHI digestion (Figure 1A(a), S1). Following lentiviral transduction, we generated OA3.Ts/Cas9 cells (Figure 1A(b)) and confirmed stable integration of Cas9 and hTERT by immunofluorescence and Western blotting (Figure 1B-C). Notably, the immortalized cells exhibited significantly enhanced proliferation compared to primary OA3.Ts cells (Figure 1D) and retained stable Cas9/hTERT expression over 107 passages, with minimal apoptosis (Figure 1A(c), S2A-E), confirming their extended replicative capacity.

**Figure 1.**
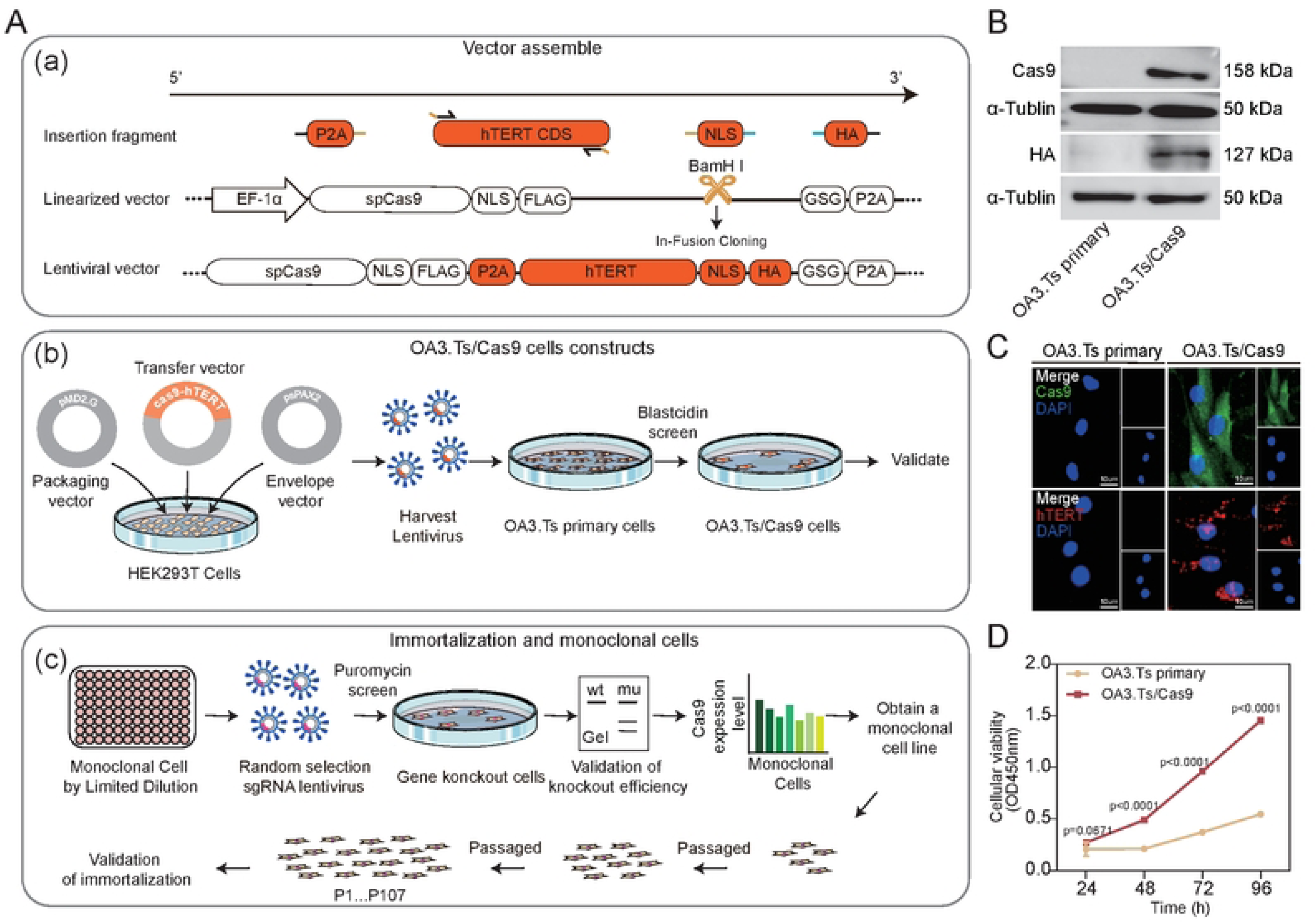
Establish immortalised OA3.Ts monoclonal cell lines stably expressing Cas9. (A) Comprehensive workflow for the generation and validation of immortalized OA3.Ts cell lines stably expressing Cas9:(a)Construction of the lentiviral expression vector Lenti-Cas9-hTERT, (b)Generation and validation of immortalized OA3.Ts cell lines stably expressing Cas9, (c)Verification and immortalization of monoclonal cells. (B) Western blot analysis of Cas9, hTERT expression in OA3.Ts primary cells and OA3.Ts/Cas9 cells. (C) Immunofluorescence staining analysis of Cas9, hTERT expression in OA3.Ts primary cells and OA3.Ts/Cas9 cells. Scale bar = 10 µm. (D) Cell proliferation activity of OA3.Ts primary cells and OA3.Ts/Cas9 cells.

To ensure functional compatibility with CRISPR screening, we isolated a monoclonal OA3.Ts/Cas9 line (clone #A) that maintained robust Cas9 expression and full susceptibility to ORFV infection (Figure S3A-D, S4A-B). To validate editing efficiency, we targeted the B4GALNT2 gene using lentiviral sgRNA delivery and observed stable CRISPR activity 6–10 days post-infection (Figure S4C-D). These results establish OA3.Ts/Cas9 as an ideal platform for genome-wide virus-host interaction studies.

### 2.2 Construction of an ovine genome-wide CRISPR/Cas9 knockout library

We developed a comprehensive ovine genome-wide CRISPR/Cas9 knockout library targeting 20,398 protein-coding genes (Figure 2A). sgRNAs were designed against all RefSeq transcript isoforms in the Ovis aries reference genome (ARS-UI_Ramb_v2.0), prioritizing 20-bp sequences with 3′-NGG PAM motifs in exonic regions. To enhance breed compatibility, we excluded sgRNAs showing sequence divergence across three major sheep breeds (Mongolian, Dairy, and Small-tailed Han sheep). Using CRISPR-offinder (v1.2), we designed 118,620 high-specificity sgRNAs, supplemented with 1,000 non-targeting negative controls (Figure S5A and Supplementary Table 1). Guide RNAs were optimized for upstream exon targeting (Figure 2B) and minimal off-target activity (Figure S5B).

**Figure 2.**
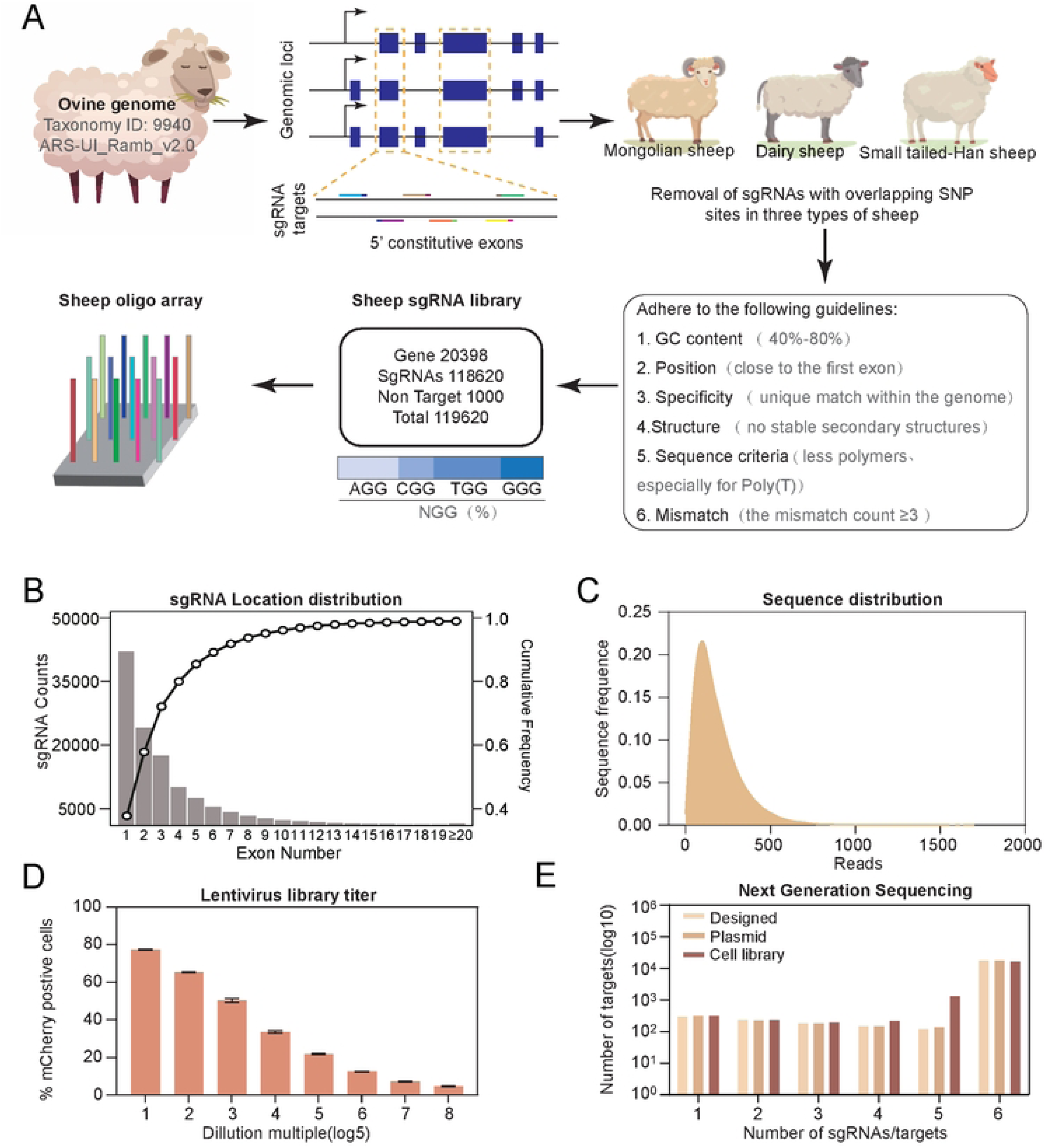
Generation of an ovine genome-wide CRISPR/Cas9 knockout library. (A) Schematic representation of the ovine genome-wide CRISPR/Cas9 knockout library design and construction workflow. Protein-coding genes from NCBI. (B) Distribution of cutting sites of all 118596 sgRNAs relative to the exon of across the genome. Bar chart showing the total number of gRNAs under the current exon. Line graph showing the proportion of frequencies under the current exon. (C) Sequence distribution of sgRNAs targeting sequences in plasmid-pooled sgRNA libraries. (D) Genome-wide lentiviral library titer assay. Characterization by flow cytometry to detect the proportion of cells with red fluorescence. All sgRNAs were inserted into the Lenti-CRISPR-puro-sgRNA-mCherry expression vector. (E) The number of sgRNAs per gene in the genome-wide CRISPR pooled sgRNA library from the designed, plasmid, or mutant cell pools. The sgRNA was designed by the CRISPR-offinder software. Plasmid, the sequencing result of the sgRNA library from plasmid pools; Cell pool, the sequencing result of the sgRNA library from sorted mutant cell populations.

Deep sequencing of PCR-amplified sgRNA constructs confirmed 99.97% (119,579/119,620) guide representation (Figure 2C, Figure S5C). Validation assays on three host genes confirmed efficient indel induction (up to 65.27% cutting efficiency) at intended loci (Figure S5D).The library demonstrated exceptional uniformity (Gini Index = 0.08234), far below the 0.2 threshold for balanced representation. Although minor abundance variations were observed, 99.76% of sgRNAs fell within a 10-fold frequency range, underscoring robust library quality.

For functional screening, we transduced OA3.Ts/Cas9 cells with the lentiviral library at a low MOI (≈0.2), achieving 20–30% transduction efficiency to ensure single-guide integration (Figure 2D, Figure S5E). The resulting cell library retained 98.61% (117,958/119,620) of sgRNAs with high uniformity (Gini Index = 0.1151; Figure 2E), establishing ovine-GeCKO as a robust resource for ovine functional genomics.

### 2.3 Genome-wide CRISPR screen identifies host factors restricting ORFV infection

To systematically identify host genes conferring resistance to ORFV infection, we performed a FACS-based genome-wide CRISPR/Cas9 knockout screen (Figure 3A). Cells from the knockout library were infected with ORFV (MOI=2), and the top 5% of cells exhibiting high viral protein expression were sorted. Genomic DNA from sorted and control cells underwent sgRNA amplification and next-generation sequencing(Figure S6). MAGeCK analysis revealed sgRNAs significantly enriched in infected cells, pinpointing antiviral host factors. The top ten most enriched candidate genes after ORFV infection were (from highest to lowest) RAD52, USP45, CSNK1G3, DNAH8, CDH13, IL22, LRP12, PEX11B, GJB6, and SLC12A5 (Figure3B and Supplementary Table 2). We observed enrichment for 0.1% Top genes involved in regulation of MAPK cascade, which were also represented in previous genome-wide screens for other viruses [34] (Figure 3C and Supplementary Table 3). To narrow the focus for further analysis, we created a score ranking of all sgRNAs for the top ten most enriched genes from the screening strategy, as well as the number of good sgRNAs determined to be statistically significant based on statistical analysis (Fig. 3D and E),we generated polyclonal knockout lines for functional validation (Figure S6A).

**Figure 3.**
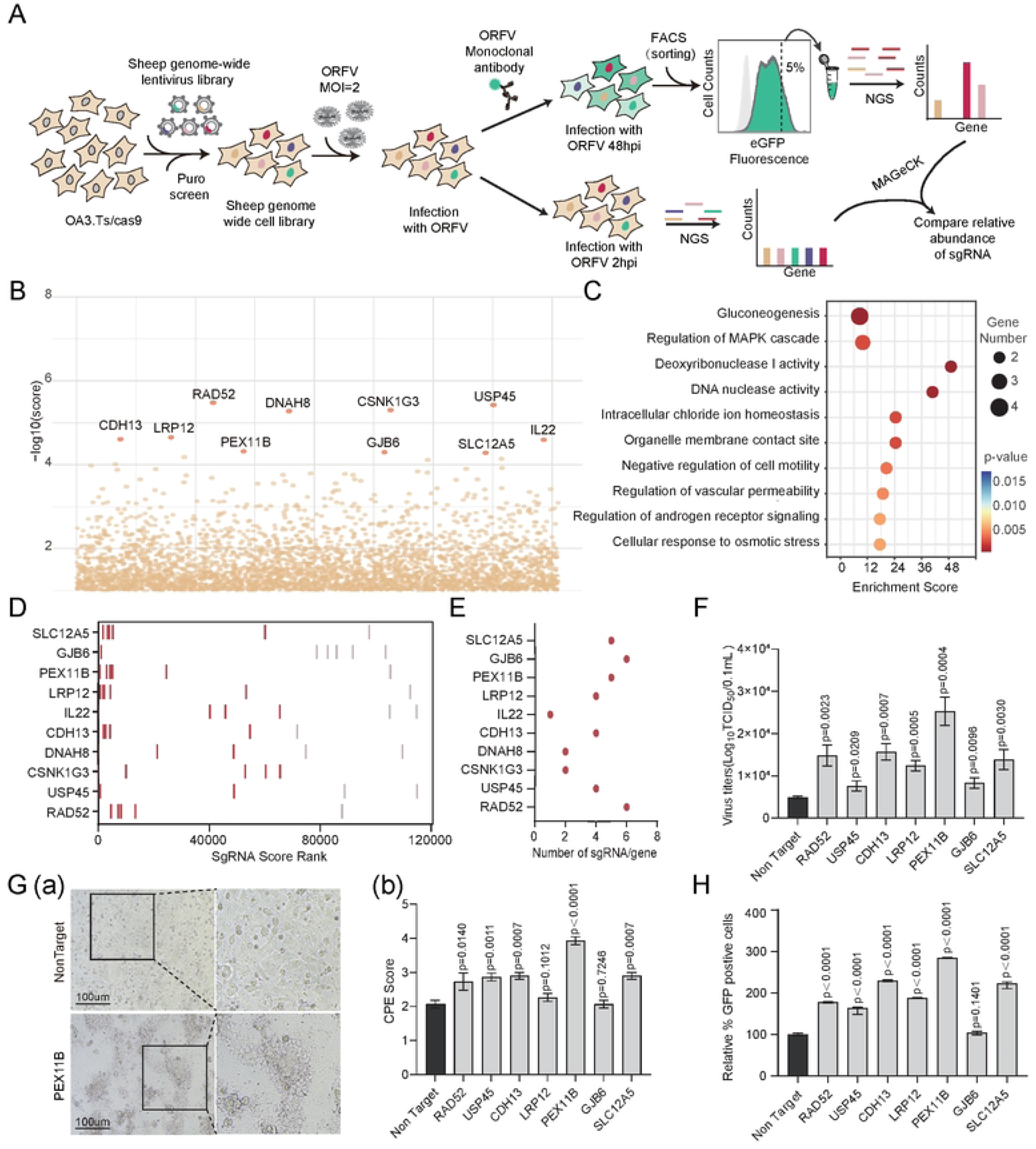
Identification of candidate genes resistant to ORFV infection via genome-wide CRISPR screen. (A) Schematic overview of FACS-based screening strategy. The sheep genome-wide CRISPR knockout library of OA3.Ts/Cas9 cells was infected with ORFV of MOI=2. After 48hpi, a FACS-based positive selection for cells was performed. Deep sequencing identified candidate genes. (B) Spatial distribution of CRISPR knockout signals on the genome for the FACS-based positive screens. The core set of the Top ten robust hits is shown in red. (C) Gene Ontology (GO) Enrichment Analysis of the Top 0.1% candidate genes. (D) Top ten candidates ranked by MAGeCK enrichment score (higher rank indicates greater enrichment). The six guides for each candidate are ranked by individual enrichment. Enriched gRNAs with Benjamini–Hochberg corrected Wald test P < 0.001 are colored in red. (E) The sgRNAs of the top ten candidates are summarised by the number of good sgRNAs in MAGeCK. (F) Seven knockout polyclonal cells were infected at MOI=0.01 ORFV. The supernatant virus titre was determined 48 hours after infection. (G) (a) Cytopathic effect (CPE) images of non-target versus KO-PEX11B cells following infection with ORFV (MOI = 2) at 36 hours post-infection. Scale bar = 100 µm. (b)The viral Cytopathic effect (CPE) scoring plot of seven knockout polyclonal cells infected with ORFV at an MOI of 2 is presented. The scoring is based on the degree of lesions: 0 = no lesions. 1: less than 25% cellular lesions. 2: 25%-50% cellular lesions. 3: 50%-75% cellular lesions. 4: more than 75% cellular lesions. (H) Flow cytometry-based relative quantitative analysis of ORF086 fluorescence intensity in seven ORFV-infected knockout polyclonal cell lines.

We next assessed ORFV infection dynamics in polyclonal knockout lines for the seven candidate genes. Knockout of all seven top candidate genes augmented ORFV infection, with PEX11B depletion showing the strongest effect (Figures 3F). In particular, ORFV infection 36 hour post infection associated with the PEX11B gene knockout showed a significant number of membrane fusion events within the cytopathic effect (CPE) (Figures 3G, S6B), corroborated by elevated infected cells and viral protein expression via immunofluorescence and flow cytometry (Figures 3H), implicating these genes as critical ORFV barriers.

### 2.4 PEX11B is a broad-spectrum gatekeeper of viral infection

Among the seven highest-ranked genes identified in the genome-scale CRISPR screen for candidates associated with ORFV infection, PEX11B, a gene implicated in peroxisome proliferation [35–37], was highlighted. Clonal PEX11B knockout cells exhibited normal proliferation (Figure S7A-B), with whole-genome sequencing confirming target specificity and complete gene ablation (Figures S7C–D), enabling mechanistic studies.

To pinpoint the infection stage modulated by PEX11B, we systematically interrogated the ORFV life cycle. Viral attachment remained unaltered in KO cells, as demonstrated by equivalent B2L envelope protein binding (Figure 4A) and consistent flow cytometry/IFA results (Figures 4B,C). Strikingly, PEX11B ablation increased viral genome internalization by 1.5 hpi (Figure 4D), while acid-bypass assays confirmed its specific role in endocytosis-dependent entry(Figure 4E). B2L mRNA levels surged in KO cells by 6 hpi (Figure 4F), collectively establishing PEX11B as a critical checkpoint for early ORFV invasion.

**Figure 4.**
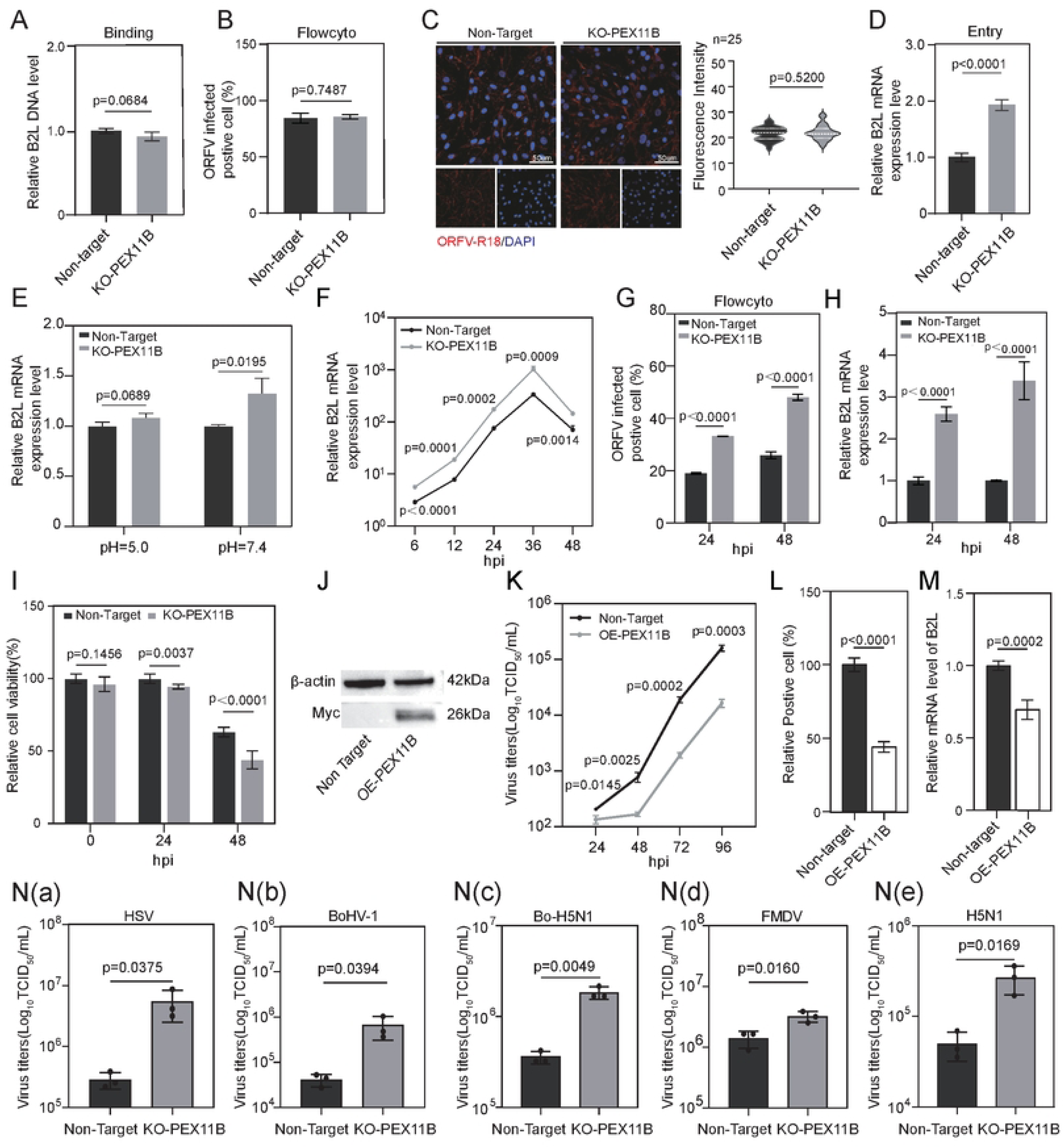
Identification of viral life-cycle defects in clonal PEX11B knockout cells. (A, B, C) Quantitative analysis of ORFV binding. Non-target versus PEX11B knockout cell lines were infected with ORFV (MOI=1), and cell surface ORFV binding was analyzed by qPCR, flow cytometry, and indirect immunofluorescence, respectively. Scale bar = 50 µm. (D) Quantitative analysis of ORFV internalization. Non-target and PEX11B knockout cell lines were infected with Orf virus (MOI = 1) for 1.5 h at 37℃, and virus internalization was analyzed by qPCR. (E) The acid bypass experiment was conducted to verify non-target and PEX11B knockout cells. (F) Quantitative analysis of B2L gene expression levels at different time points after ORFV infection. (G) Flow cytometric analysis of infected-positive cell proportions in Non-target and PEX11B knockout (KO-PEX11B) cells at 24 and 48 hours post-infection (hpi). Unpaired t-test, p < 0.0001 for both time points. (H) Relative ORFV mRNA levels in Non-target and KO-PEX11B cells at 24 and 48 hpi (normalized to an endogenous control). Unpaired t-test, p < 0.0001 for both time points. (I) Relative cell viability of non-target and KO-PEX11B cells at 0, 24, and 48 hpi after ORFV infection at MOI = 5. (J) Western blot analysis of Myc-flag expression in non-target cells and PEX11B overexpression cells. (K) Non-Target and PEX11B overexpression lines were challenged with ORFV (MOI=0.001), and the viral titers were measured at the indicated time post-infection. (L 、 M) Non-Target and PEX11B overexpression cell lines were challenged with ORFV at 24 hours post-nfection. Fluorescence of the virus on the cell was shown in (L), and relative expression of ORF viruses was measured in (M). (N) Virus titer assays for Non-target and PEX11B knockout cells infected with some virus.(a)HSV (MOI = 0.1, 24 hpi).(b) BoHV-1 (MOI = 1, 24 hpi).(c) Bo-H5N1 (MOI = 0.25, 24 hpi).(d) VN/H5N1 (MOI = 0.25, 24 hpi).

Post-entry, PEX11B KO increased infected cells (24-48 hpi; MOI=5) (Figure 4G) and viral mRNA (Figure 4H), exacerbating cell death (Figure 4I). In contrast, PEX11B overexpression (OE) suppressed ORFV titers, infection rates, and B2L expression (Figures 4J-M), confirming dosage-sensitive antiviral activity.

To determine whether PEX11B’s antiviral function extends beyond ORFV, we challenged KO cells with phylogenetically distinct viruses: the DNA viruses HSV-1 and BoHV-1(Bovine herpesvirus 1), and RNA viruses BoH5N1(Bovine H5N1) and VN/H5N1 (Influenza A). All viruses exhibited significantly enhanced replication in KO cells (Figure 4N), demonstrating PEX11B’s capacity to restrict diverse viral families.

Our systematic analyses position PEX11B as a central hub of an underappreciated host defense pathway that limits infection by multiple viral families. The conservation of this restriction across DNA and RNA viruses suggests PEX11B may represent an evolutionarily ancient antiviral mechanism co-opted by diverse viral pathogens.

### 2.5 Peroxisome biogenesis is a proviral factor for ORFV replication

To delineate the role of peroxisomes in ORFV infection, we first analyzed the spatiotemporal relationship between viral replication and peroxisome dynamics via immunofluorescence microscopy. Infection with ORFV induced a striking 1.98-fold increase in peroxisome abundance by 72 hpi (Figure 5A), suggesting a functional link between peroxisomal proliferation and viral replication.

**Figure 5.**
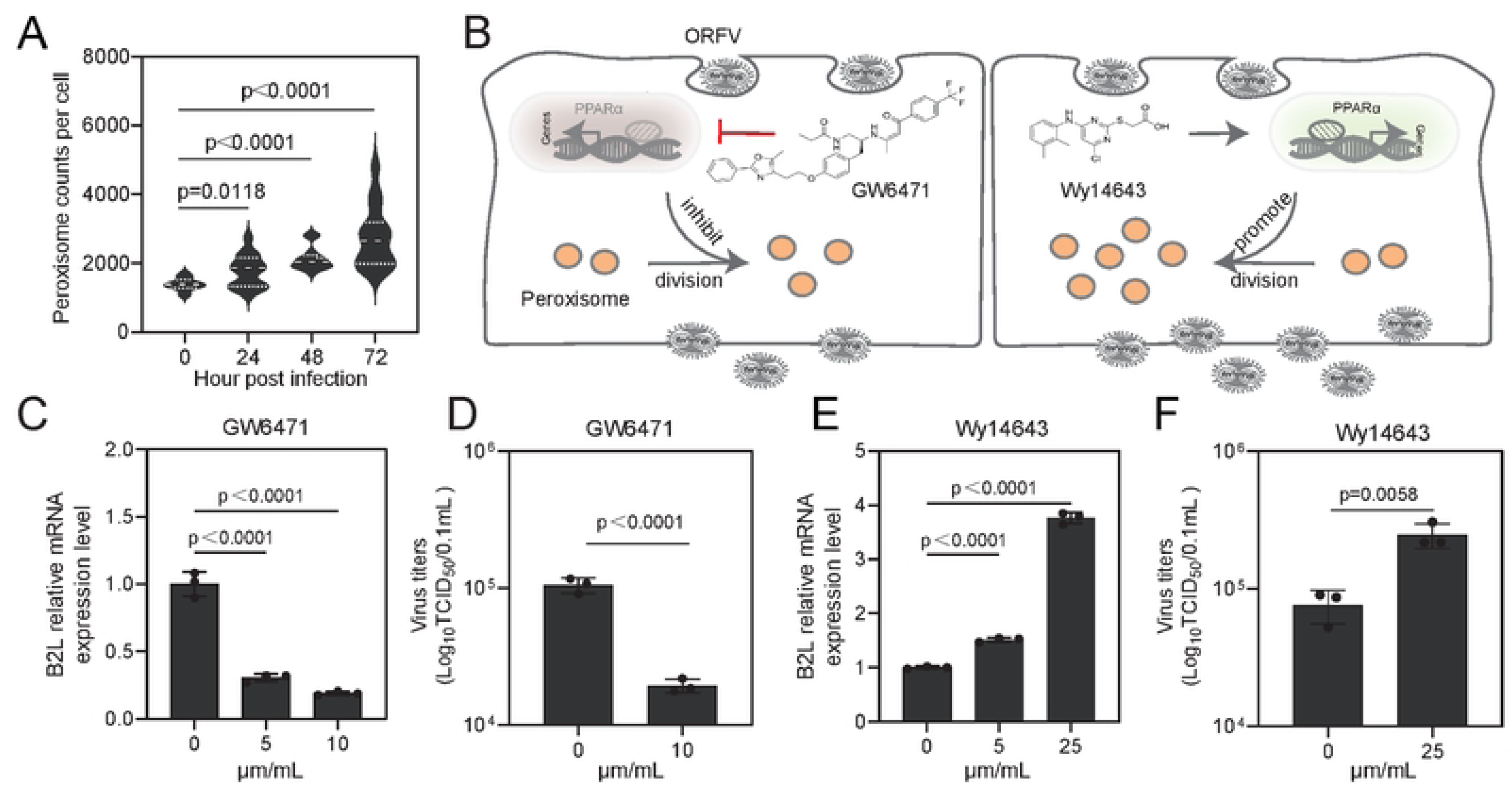
Antiviral function of PEX11B mediated by the action of peroxisomes. (A) Peroxisome numbers in ORFV-infected OA3.Ts/Cas9 cells. N > 13 cells. (B) Schematic illustration depicting that ORFV infection drives peroxisome fission via the PPARα pathway, an effect that can be pharmacologically suppressed by GW6471 or potentiated by Wy14643. (C、D)ORFV infectious B2L gene mRNA expression level and virus titer produced from infected OA3.Ts/Cas9 cells following GW6471 treatment to inhibit peroxisome biogenesis. (E、F)ORFV infectious B2L gene mRNA expression level and virus titer produced from infected OA3.Ts/Cas9 cells following Wy14643 treatment to induce peroxisome biogenesis.

To further prove this relationship, we pharmacologically modulated peroxisome biogenesis in OA3.Ts/Cas9 cells using GW6471 (PPARα antagonist, 5 - 10 μg/mL) and Wy14643 (PPARα agonist, 5 - 25 μM) (Figure 5B) [38, 39].Pharmacological inhibition of PPAR α with GW6471 induced a dose-dependent suppression of viral replication, reducing B2L transcript levels to 19 - 30 % and viral titers to ∼18 % relative to vehicle-treated controls (Fig. 5C-D and S8A).In contrast, PPARαactivation with Wy14643 enhanced viral replication, increasing B2L expression by 1.5 - to 3.8-fold and viral titer by 3.2 - fold (Figure 5E-F, S8B). Taken together, these findings establish peroxisome abundance as a critical determinant of ORFV replication efficiency, where increased peroxisomal biogenesis positively correlates with enhanced viral propagation.

### 2.6 PEX11B maintains peroxisomal membrane homeostasis to restrict ORFV infection

PEX11B regulates peroxisomal growth and division by coordinating membrane remodeling and fission machinery assembly [40]. To dissect its antiviral role, we made PEX11B mutants missing important sections (Δ159-183, Δ185-201, and Δ210-259) (Figure 6A, S9A). Intriguingly, while the Δ159-183 and Δ210-259 mutants reduced peroxisome abundance to PEX11B knockout levels (Figure 6B), both mutants lost antiviral activity, as evidenced by enhanced ORFV replication (Figure 6C-D, S9B). PEX11B therefore serves as a key controller of peroxisome-based defense, regardless of peroxisome count.

**Figure 6.**
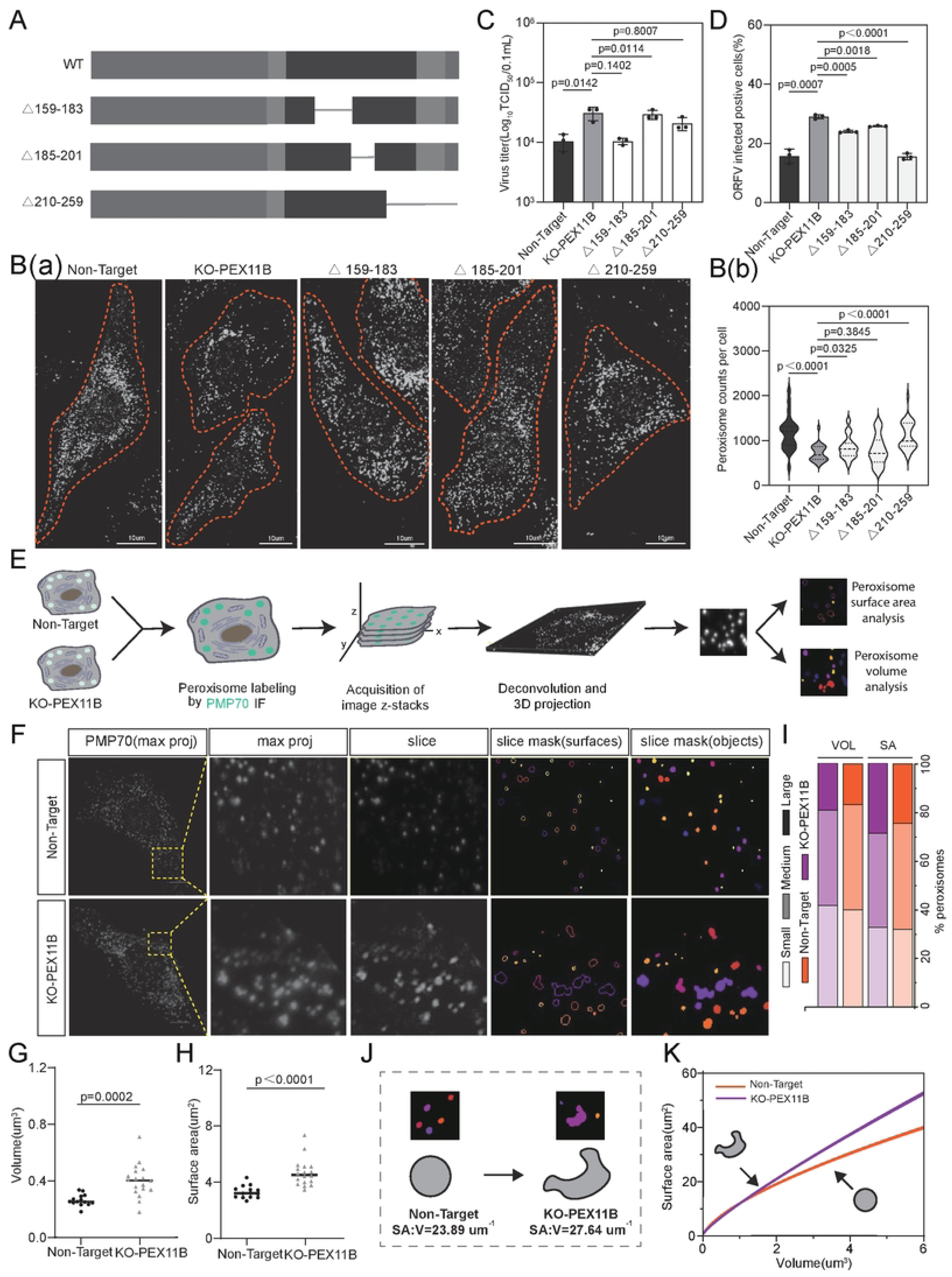
PEX11B knockout alters peroxisomal Membrane-to-Lumen Ratio to enhance ORFV Infection. (A)Schematic of wild-type (WT) PEX11B and the PEX11B mutants △210-259,△159-183,△185-201. (B)(a)Representative images of peroxisome morphology (anti-PMP70) in non-target, PEX11B knockout cells, and PEX11B mutants △210-259,△159-183,△185-201 cells. (b)Statistical summary of the number of peroxisomes quantified. Non-Target N=26793,PEX11B KO cells N=19407,PEX11B mutants(△210-259) N=33297,PEX11B mutants(△159-183) N=27353,PEX11B mutants(△185-201) N=24201. Scale bar = 10 µm. (C) Analysis of the ORFV (MOI=0.01,48h) virus titer through TCID50. (D) Analysis of the proportion of positive cells through flow cytometry. Data are presented as the mean ± SD of triplicate measurements from a single experiment, with consistent results observed in at least three independent replicates. (E) Workflow for the analysis of peroxisome surface area and volume. (F) Non-target and KO-PEX11B cells labeled with anti-PMP70. ROI squares (left) show z-stack maximum projection, two-slice mask (surfaces and objects), and masks of peroxisomes from object analysis. Small peroxisomes (arrows) and enlarged peroxisomes (arrowheads) are indicated. Scale bar = 10 µm. (G) Average volume peroxisome morphology and (H)average surface area per cell in NonTarget and KO-PEX11B conditions. N = 14 cells in NonTarget; N = 18 in PEX11B KO Mock. (I) Analysis of peroxisome size distribution in non-target and KO-PEX11B cells. Peroxisome surface area (SA) and volume (VOL) were quantified and binned into small (SA, < 0.9 mm2; VOL, < 0.06 mm3), medium (SA, 1.1–3.3 mm2; VOL, 0.07–0.34 mm3), and large (SA, > 3.3 mm2; VOL, > 0.34 mm3) categories.N =8138 peroxisomes in mock; N = 7882 peroxisomes in ORFV. (J) Peroxisome morphology changes. Examples of peroxisome masks from 3D analysis are shown, with SA and VOL values indicated. (K) Surface area-to-volume ratio (SA: V) regression curves from data in (G and H), plotted in the physiologically relevant range of observed mock peroxisome volumes (1–6 mm3). Significance of *p* < 0.001 was determined by the chi-squared test for (I). Significance was determined by Student’s t-test for (G and H), marked with p.

To further understand these effects, we used 3D imaging and fluorescent microscopy to examine peroxisome shape (Figure 6E). In healthy cells, peroxisomes were evenly round (Figure 6F), while KO-PEX11B cells showed larger, oddly shaped peroxisomes (Figure 6F, S9C). Shape analysis showed an uneven size distribution in KO-PEX11B cells (Figure 6G-I). Importantly, KO-PEX11B peroxisomes had a much higher surface area-to-volume (SA:V) ratio (Figure 6J), and further modeling found that as volume increased, so did SA:V (Figure 6K, S9D), resulting in smaller inner space and more surrounding membrane—a structure that may help viruses take advantage.

### 2.7 PEX11B deficiency reprograms lipid metabolism to elevate membrane fluidity

Given peroxisomes’ central role in lipid metabolism[41, 42], we performed LC-MS lipidomics on KO-PEX11B cells and identified 624 dysregulated lipids (445 upregulated, 179 downregulated; p < 0.01). There were pronounced accumulations of triglycerides (TG) and phosphatidylcholine (PC). In contrast, we also observed depletions of phosphatidylethanolamine (PE) and sphingomyelin (SM) (Figures 7A-C, S10A-C). Notably, PEX11B ablation led to increased ratios of PC:PE and PC/sphingomyelin (SM) (Figures 7D-E, S10D-E), a perturbation known to reduce lipid packing density and increase membrane permeability (41-43). Furthermore, PEX11B-KO cells exhibited a higher total unsaturated-to-saturated and mono- and diunsaturated-to-saturated fatty acid ratio (Figures 7F-G), suggesting enhanced desaturation that promotes membrane fluidity by increasing acyl chain disorder. Finally, heatmap analysis implicated PEX11B loss in activating the glycerophospholipid/sphingolipid metabolic pathway (Figure 7H), thus directly linking peroxisomal dysfunction to altered membrane properties.

**Figure 7.**
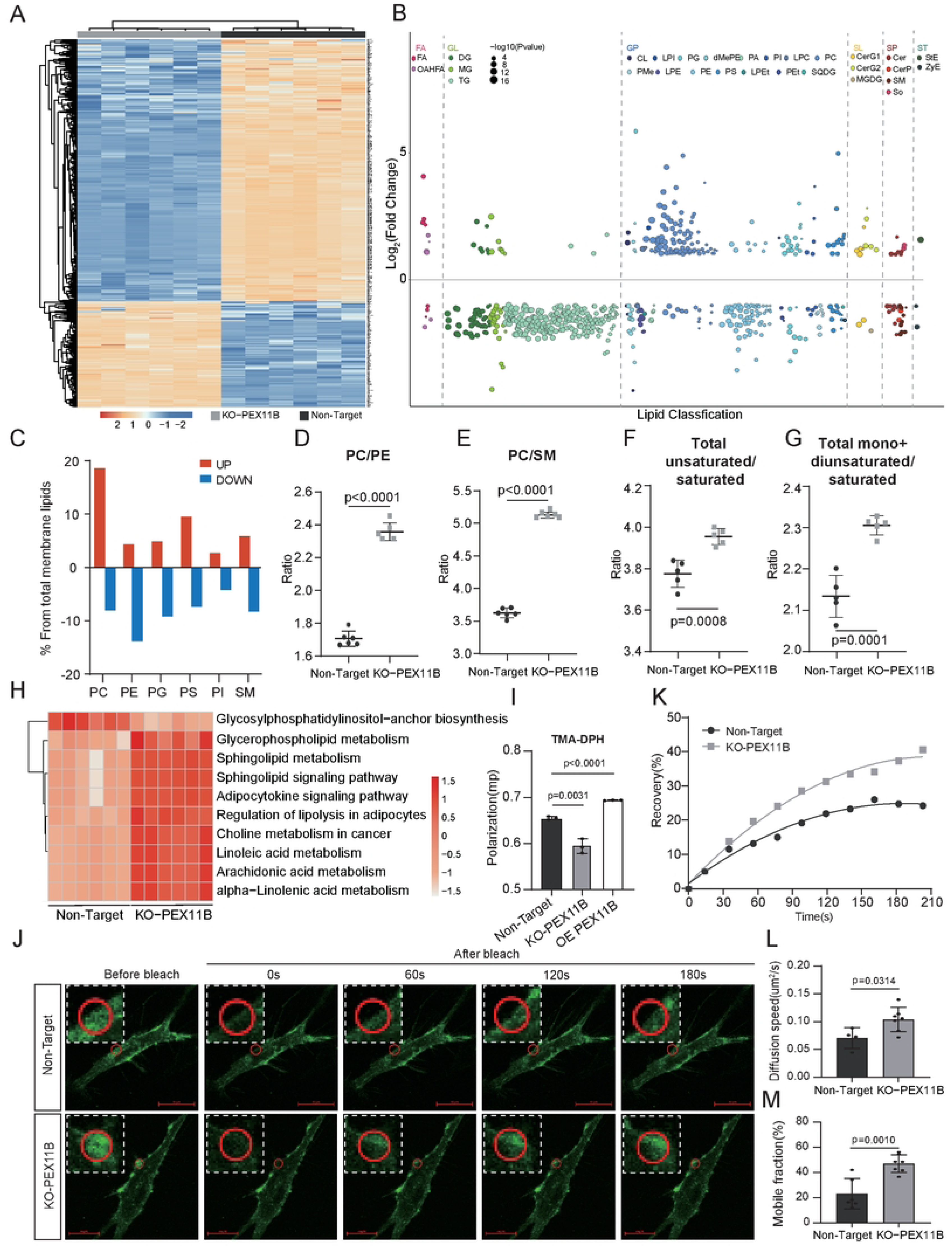
Knockout of PEX11B promotes cell membrane fluidity. (A) Heatmap showing the lipidomic analysis of non-target and KO-PEX11B cells. Each rectangle represents an ion feature colored by its normalized intensity scale from red (decreased level) to yellow (increased level). The dendrogram at the top was constructed based on the lipid intensity (similarity measure using Euclidean distance and the Ward clustering algorithm). (B) Bubble plots for correlation analysis of differential lipid classification. (C) Changes in the composition of cell-associated membrane lipids in non-target cells and KO-PEX11B cells. (D 、 E) Scatter plot showing the ratios of concentrations of total phosphatidylcholine versus total phosphatidylethanolamine lipids and total phosphatidylcholine versus total sphingomyelin lipids in the non-target cells and KO-PEX11B cells. (F、G) Scatter plot showing the ratios of concentrations of total unsaturated versus total saturated lipids and total mono+diunsaturated versus total saturated lipids in the non-target cells and KO-PEX11B cells. (H) KEGG pathway analysis of bulk LC-MS data with averaged lipid expression shows the dynamic changes of lipid metabolism pathways across non-target cells and KO-PEX11B cells. (I) Fluorescence polarization (P) value changes of TMA-DPH-stained non-target cells, KO-PEX11B cells, and PEX11B overexpression cells. P was measured at an excitation wavelength of 355 nm and an emission wavelength of 430 nm. Data were collected from three parallel experiments. P is depicted as means ± SD, n = 3. (J) Fluorescence recovery images of non-target cells (Top) and KO-PEX11B cells (Bottom) after photobleaching upon various treatments. The radius of the selected region is 2.5 μm for all images. Scale bar = 20 μm. (K) Plots of fluorescence intensity of the non-target cells and KO-PEX11B cells of the marked area in (J) versus time after photobleaching. (L) Diffusion speed and (M) Mobile fraction of FITC-labeled cell membrane obtained from plots of normalized FRAP data. The means and SDs were from at least three different samples.

To test the hypothesis that PEX11B knockout increases membrane fluidity through altered phospholipid balance and fatty acid unsaturation, we performed biophysical assays. TMA-DPH fluorescence polarization, which measures membrane rigidity by tracking fluorophore rotation, revealed significantly reduced anisotropy (*P* = 0.595 ± 0.002) in KO-PEX11B cells compared to non-target (0.653 ± 0.001) and OE-PEX11B cells (0.693 ± 0.001), confirming enhanced fluidity (Figure 7I). In line with these results, fluorescence recovery after photobleaching (FRAP) assays—used to assess lateral membrane diffusion—using DiOC18 demonstrated accelerated recovery kinetics in KO-PEX11B cells, with a 5% higher diffusion coefficient (0.068 ± 0.025 μm²/s) and an increased FRAP ratio (0-40%) relative to controls (0-24.2%, 0.104 ± 0.035 μm²/s) (Figures 7J-L). Additionally, the mobile fraction (Mf), an indicator of the proportion of lipids able to move freely within the membrane, exhibited significant differences between groups (p < 0.005, Figure 7M). Together, these findings establish PEX11B as a modulator of membrane fluidity, offering mechanistic insights into peroxisome-virus interactions.

### 2.8 Increased lipid membrane fluidity promotes ORFV infection

To systematically assess the impact of host membrane fluidity on viral infection, we infected isogenic cell lines (KO-PEX11B, Non-target, and OE-PEX11B) with ORFV. Because these cell lines exhibited graded membrane rigidity, we quantified infection efficiency relative to TMA-DPH fluorescence polarization, a marker of reduced membrane fluidity (Figure 8A). Notably, KO-PEX11B cells with elevated fluidity supported significantly higher ORFV entry (Figure 4D), suggesting a fluidity-dependent penetration mechanism. To dissect this further, we performed time-resolved confocal imaging of R18-labeled ORFV, revealing accelerated endosomal transit in KO cells: viral particles showed enhanced Rab5+ early endosome colocalization by 10 min post-infection (Figure 8B), rapid progression to Rab7+ late endosomes by 20 min (Figure 8C), and premature Lamp1+ lysosomal accumulation by 30 min (Figure 8D). These timepoints coincide precisely with the viral uncoating phase[43–45]. Further supporting this, lysosomal activity assays confirmed heightened endolysosomal function in KO cells (Figure 8E). Collectively, these results establish membrane fluidity as a critical positive regulator of ORFV entry by accelerating vesicle maturation.

**Figure 8.**
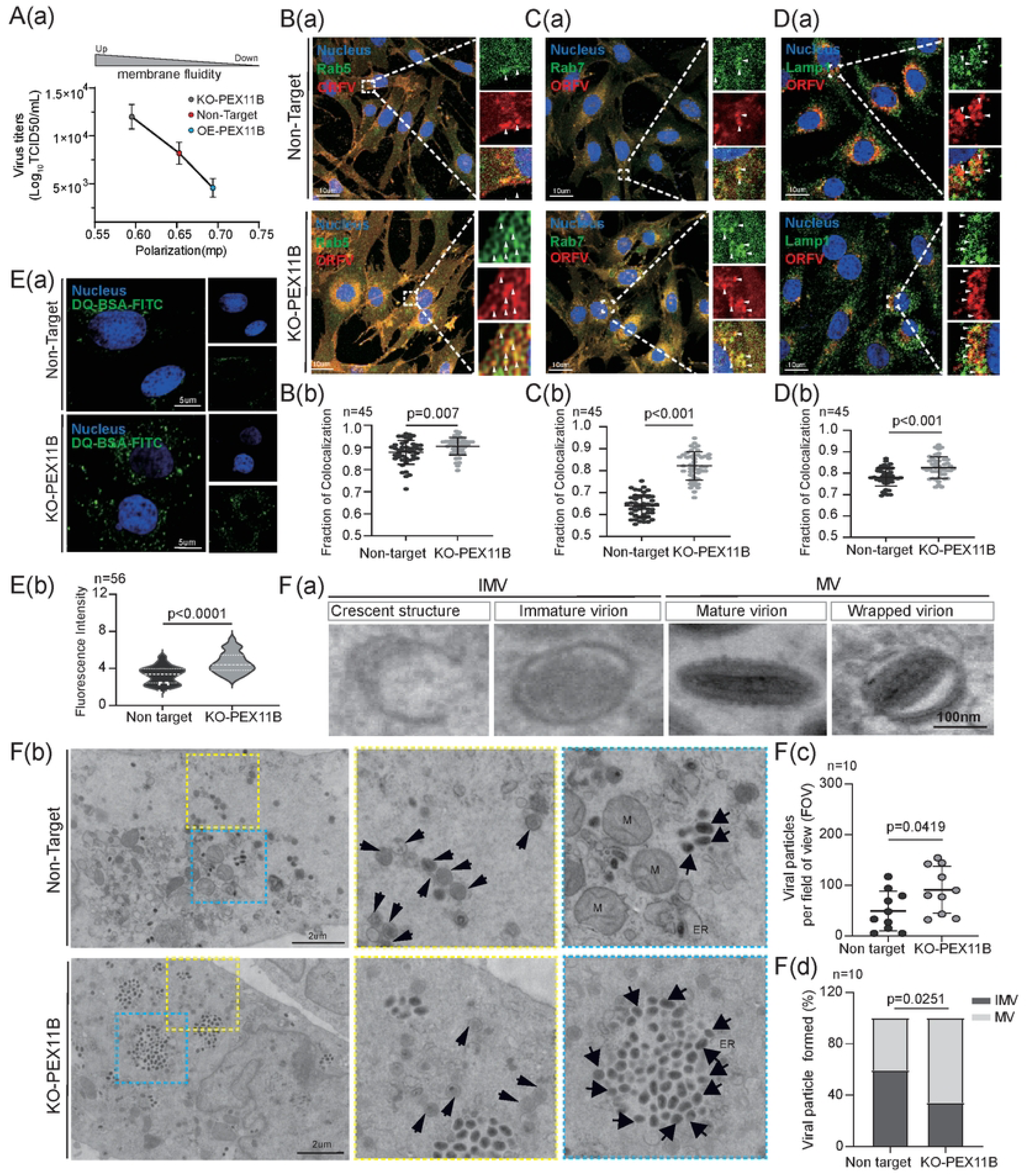
Increased lipid membrane fluidity promotes ORFV replication. (A) Cell lines with graded membrane rigidity (KO-PEX11B, non-target, and OE-PEX11B) were infected with ORFV, and the correlation between infection efficiency and TMA-DPH fluorescence polarization was analyzed. Data show an inverse correlation between viral infection efficiency and TMA-DPH fluorescence polarization. (B)(a)Confocal microscopy analysis of co-localization of ORFV and endosomes in non-target versus PEX11B knockout cell lines, 10 min (GFP-Rab5 early endosomes) after infection.Blue, Hoechst 33258; red, R18-labeled ORFV; green GFP, Rab5 early endosomes.(b) Histograms of Rab5 early endosomes and virus co-localization analysis, cell n=45. (C) (a)Confocal microscopy analysis of co-localization of ORFV and endosomes in non-target versus PEX11B knockout cell lines, 20 min (GFP-Rab7 late endosomes) after infection. Blue, Hoechst 33258; red, R18-labeled ORFV; green GFP, Rab7 late endosomes.(b) Histograms of Rab7 late endosomes and virus co-localization analysis, cell n=45. (D) (a)Confocal microscopy analysis of co-localization of ORFV and endosomes in non-target versus PEX11B knockout cell lines 30 min (GFP-Lamp1 lysosomes) after infection. (b) Histograms of Lamp1 lysosomes and virus co-localization analysis, cell n=45. (E) Confocal microscopy analysis of non-target, PEX11B knockout cells using DQ-green bovine serum albumin (BSA), Scale bar = 10 µm. The violin plot depicts the relative fluorescence intensity for different cells. Colocalization analyses are consistent with the previous description, n=56. Scale bar = 5 µm. (F) (a) Representative TEM images illustrating sequential stages of ORFV assembly: crescent structure, immature virion, mature virion, and wrapped virion. Scale bar, 100 nm. (b) TEM micrographs of ORFV-infected non-target and KO-PEX11B cells. Arrows indicate viral particles; yellow and blue boxes denote regions magnified in adjacent panels. Small arrow indicates immature virion; large arrow indicates mature virion. Scale bar, 2 μm. (c) Quantification of viral particles per field of view (FOV) in Non-target and KO-PEX11B cells (n = 10 FOVs). Data are presented as mean ± SEM; unpaired t-test, p = 0.0419. (d) The proportion of Intracellular Mature Virions (IMV) and Mature Virions (MV) in non-target cells and KO-PEX11B cells. Unpaired t-test, p = 0.0251.

Transmission electron microscopy of KO-PEX11B and non-target cells revealed conserved stages of ORFV morphogenesis (Figure 8F(a)). Moreover, KO-PEX11B cells exhibited a marked increase in viral particle density (Figure 8F(b)). Consistent with this, quantitative morphometric analysis showed more mature, envelope-intact virions in KO-PEX11B populations (Figure 8F(c,d)). Together, these findings directly link membrane fluidity to assembly fidelity. Thus, these data establish that suppression of fluidity by PEX11B acts as an innate barrier to ORFV infection, impacting both viral entry via endosomal trafficking and late-stage virion production.

## 3 Discussion

In this study, we performed the first ovine genome-wide CRISPR/Cas9 knockout screen in primary OA3.Ts cells. This allowed us to systematically identify host factors that regulate ORFV (Orf virus) infection. Our screen identified PEX11B as a critical antiviral regulator (Figure 9). When PEX11B was knocked out, ORFV replication increased. This was shown by more syncytia at 36 hours post-infection and a 4.23-fold increase in viral titer at 48 hours. We also observed peroxisomal remodeling: fewer peroxisomes, but greater peroxisome volume and surface area. This led to a 15.7% rise in membrane-to-lumen ratio (from 23.89 to 27.64; Figure 6). These structural changes matched with higher lipid metabolic efficiency, as shown by LC-MS lipidomics (liquid chromatography-mass spectrometry), altered phosphatidylcholine/phosphatidylethanolamine and sphingomyelin ratios, increased membrane fluidity (measured by TMA-DPH and FRAP assays), and faster viral infection.

**Figure 9.**
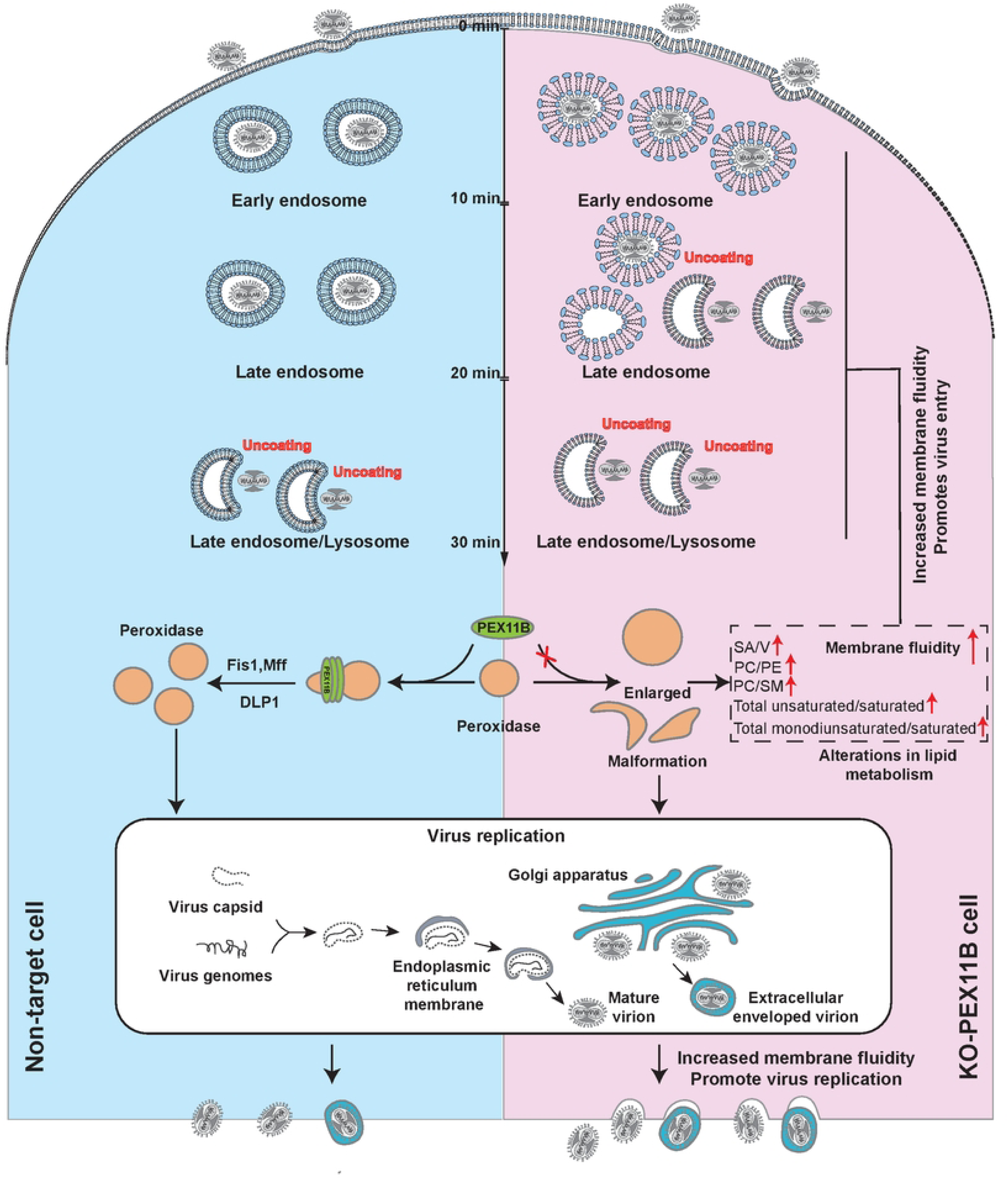
A model illustrating the roles of PEX11B in the ORFV replication cycle. This schematic outlines the role of PEX11B in regulating peroxisome dynamics and downstream ORFV infection processes. In non-target cells, PEX11B mediates proper peroxisome fission through interactions with Fis1, Mff, and DLP1, maintaining normal peroxisome morphology. In PEX11B-knockout cells, impaired peroxisome fission leads to enlarged and malformed peroxisomes, which induce alterations in lipid metabolism (e.g., changes in SA/V, PC/PE, PC/SM, and unsaturated/saturated lipid ratios). These metabolic changes first enhance lipid mobility, which in turn increases membrane fluidity and alters lipid proportions. Collectively, these effects promote key ORFV biological processes: enhanced ORFV entry (including membrane fusion), accelerated ORFV trafficking (from early to late endosomes/lysosomes with uncoating), and facilitated ORFV replication (encompassing genome replication, capsid assembly, maturation via the endoplasmic reticulum and Golgi apparatus, and release of enveloped ORFV virions). Thus, PEX11B deficiency disrupts peroxisome homeostasis, triggering a cascade of lipid-related changes that drive augmented ORFV infection.

Pooled CRISPR screening enables systematic interrogation of thousands of genetic perturbations in a single experiment. However, successful implementation requires a well-characterized, optimized cellular model [46]., with critical considerations including genetic background, growth rate, and transduction efficiency [47]. Here, we addressed these challenges by generating a Cas9-hTERT-fused OA3.Ts primary cell line, which retained primary-cell characteristics while achieving immortalization and high transduction efficiency. This breakthrough enabled the first genome-wide CRISPR/Cas9 screen in ovine cells.

To ensure robust screening, we employed low MOI (∼0.2) to limit cells to single sgRNA integrations and maintained 1,000x sgRNA coverage, enhancing reproducibility and sensitivity for detecting changes in sgRNA abundance [48]. Library diversity is inherently biased by sgRNA representation, necessitating large cell populations to capture underrepresented guides. Our approach optimized these parameters, ensuring reliable hit identification.

Conventional CRISPR screens for viral host factors often rely on cell survival or death as endpoints, limiting discovery to genes essential for viral replication while missing inhibitory factors. To address this, we employed ORFV antibody-coupled fluorescence to enrich a GFP-high population (top 5%), identifying gene knockouts that potentially enhanced infection. This approach revealed multiple novel ORFV-associated host factors, including RAD52, USP45, CSNK1G3, DNAH8, CDH13, IL22, LRP12, GJB6, and SLC12A5,which are worthy of further study.We discovered these genes involved in diverse biological processes, including gluconeogenesis, regulation of MAPK cascade, deoxyribonuclease I activity, DNA nuclease activity, intracellular chlorideion homeostasis, organelle membrane contact site, etc. Knockout validation confirmed the functional relevance of these candidates, supporting the screen’s reliability.

PEX11B, a peroxisomal membrane protein regulating fission by modulating membrane dynamics [49], exhibits an antiviral role whose mechanisms were previously unclear. We found that PEX11B knockout reduced peroxisome number but increased volume and surface area, yielding a 15.7% higher membrane-to-lumen ratio (Figure 6), suggesting morphological restructuring (e.g., spherical to tubular/dumbbell shapes) that expands membrane contact sites. Although unresolved by high-resolution imaging, the enlarged surface area likely facilitates (1)greater transmembrane transport (e.g., fatty acid uptake for viral lipid synthesis), (2) enhanced lipid metabolic flux (supplying phospholipids for viral envelopes)[50–52], and (3) improved membrane fluidity (accelerating viral entry/egress) [53]. PEX11B deficiency also disrupted lipid homeostasis, elevating PC/PE and PC/SM ratios that are crucial for lipid raft signaling and may thereby influence viral entry, glycoprotein trafficking, or assembly. While these findings align with peroxisomal lipids’ pro-viral roles[54, 55], the exact interplay between PEX11B, membrane dynamics, and ORFV restriction requires further investigation.

By establishing a genome-wide CRISPR screen in primary ovine cells, we identified PEX11B as a broad-spectrum gatekeeper of viral infection and elucidated its role in peroxisome-mediated antiviral defense. Our findings expand the understanding of peroxisomal plasticity in viral pathogenesis and highlight lipid metabolism as a critical regulatory node. The screen’s top hits, including unvalidated candidates, present promising avenues for dissecting ORFV pathogenesis and broader ovine infectious diseases. Moreover, the methodological pipeline developed here provides a scalable platform for investigating host-pathogen interactions in livestock, with potential applications in vaccine development and antiviral therapies.

## 4 Materials and methods

### 4.1 Cell culture and virus

Ovine Fetal Testis cell lines and HEK293T were cultured in high-glucose DMEM (Gibco) with 10% fetal bovine serum, 1% MEM non-essential amino acids, 100 units/ml penicillin, and 100 mg/ml streptomycin (all Gibco). Cells were incubated at 37 °C and 5% CO_2_.

### 4.2 Plasmids

pMD2.G (#12259), pSPAX2 (#12260), Lenti-guide-puro (#52963), Lenti-Blast-Cas9 (#52962) were purchased from Addgene. pMD19-T Vector (#6013) were purchased from Takara.PCDNA3.1、pLV3-CMV-MCS-3×Myc-Neo、plasmid DNA was transfected into the indicated cells using Lipofectamine® LTX Transfection Reagent (Invitrogen, Thermo Fisher Scientific, USA).

### 4.3 Construction pLV-cas9-hTERT plasmid

According to the hTERT gene sequence in the NCBI database (NCBI entry number: NM_198251.2), synthetic fragment A (produced by Shanghai Shenggong Technology Company) and fragment B (produced by Beijing Genomics Institution). Aim at fragment A and fragment B, using four pairs of specific primers:

hTERT-F1: 5’-CAAAGACGATGACGATAAGGGATCCGGCGCTACT-3’

hTERT-R1: 5’-CACGCACACCAGGCACT-3’; hTERT-F2: 5’-GTTTGTCCAGGATGGTCTTGA-3’, hTERT-R2: 5’-AGTTTGTTGCGCCGGAT-3’;

hTERT-F3: 5’-CAAAGACGATGACGATAAGGGATCCGGCGCTACT-3’, hTERT-R3: 5’-CGACGTAGTCCATGTTCACAATC-3’;

hTERT-F4: 5’-GAGCTGCTCAGGTCTTTCTTT-3’, hTERT-R4: 5’-AGTTTGTTGCGCCGGAT-3’.

Fragment A served as the template for PCR amplification, and fragments 1 (259 bp) and 2 (111 bp) (111 bp) were obtained, respectively. Fragment 3 (2104 bp) was generated through PCR amplification using fragment B and fragment one as templates. Additionally, fragment 4 (1828 bp) was obtained by PCR amplification using fragment B and fragment two as templates.The PCR amplification system is: 2×PrimerSTAR GXL Buffer 10μL, dNTP Mixture (2.5mM) 4μL, Primer-F 0.5μL, Primer-R 0.5μL, Template 1μL (100ng), PrimerSTAR GXL DNA Polymerase 1μL, ddH2O 33μL.Amplified fragment three and fragment four and pLenticas9-Blast plasmid (Addgen, 52962) were digested with BamHI and then ligated using Infusion DNA ligase (Takara, China). Then the conjugated product was transformed by the receptive Escherichia coli. Colony PCR screening for positive Pcas9-hTERT clones using specific primers (TF/TR). Plasmid constructs were isolated from some positive clones by Plasmid Miniprep Kit (QIAGEN, Germany) after being grown in LB/ampicillin medium. Successful cloning of these plasmid constructs was confirmed by digestion with restriction enzymes and by sequencing.

### 4.4 Construction of OA3.Ts/Cas9 cell line

To construct the immortalized OA3.Ts cell line overexpressing Cas9 (OA3.Ts/Cas9), the Cas9 expression cassette was inserted into the human telomerase reverse transcriptase (hTERT) gene. The Cas9 expression cassette was amplified from Lenti-Blast-Cas9 (#52962) and linearised through BamHI digestion, followed by seamless cloning with the purified hTERT fragment using Infusion cloning. Subsequently, the construct was packaged into lentivirus using HEK293T cells and transfected into primary OA3.Ts cells. After 14 days of selection with blasticidin (Coolaber, CB2812) 5ug/mL, single clones overexpressing Cas9 were obtained through limiting dilution and confirmed by western blot analysis for Cas9 and hTERT expression. The established cell line was continuously passaged for over one hundred generations to assess its potential for immortalization.

### 4.5 Ovine genome-wide sgRNA library design

An ovine genome-wide CRISPR/Cas9 sgRNA knockout library consisting of 118,620 CRISPR sequences targeting 20,398 protein-coding genes, along with 1,000 non-target CRISPR sequences, was synthesized as 80-mer single-stranded oligos in the following format: TATATCTTGAGGAAAGGACGAAACACCG-N20-GTTTTAGAGCTAGAAATAGCAAGTTAA (N20:variable 20 bp CRISPR sequences). The oligos were then converted to double-stranded DNA (dsDNA) through PCR and cloned into SYN004-pKLV-U6gRNA(BbsI)-PGKpuro2A-mcherry by BbsI digestion and T4 ligation to make the ovine genome-wide CRISPR/Cas9 knockout sgRNA library. All CRISPR sequences were searched with an NGG protospacer adjacent motif (PAM) at the 3′ end of the exon regions of all protein-coding genes in the ovine genome (Taxonomy ID: 9940, reference genome: ARS-UI_Ramb_v2.0). Additionally, please adhere to the following guidelines. Ensure that the GC content is approximately between 40%-80%. 2. Preferably, select the first exon region for cleavage sites. 3. Analyze the specificity of the sgRNA; the target sequence should have a unique match within the genome. 4. The sgRNA sequence should not contain stable secondary structures, such as hairpin structures. 5. The designed gRNA sequence (excluding the PAM) should not contain four or more consecutive thymine (T) nucleotides. 6. Consider the efficiency score predicted for the gRNA sequence. 7. The potential off-target or mismatch count for the gRNA sequence should be three or more. To expand the application of the library across different sheep breeds, the library simultaneously selected the gene sequences of three sheep breeds, namely the Hu sheep, dairy sheep, and Mongolian sheep, as references, and removed the sgRNAs that had differences among the three breeds.

### 4.6 Construction of the ovine CRISPR/Cas9 sgRNA plasmid library

The PCR reaction was conducted in a Veriti™ 96-Well Thermal Cycler (Thermo Fisher Scientific) with 14 cycles. A total of 35 PCR reactions were performed, each containing 20 ng of oligo pool per 50 μL reaction volume. The PCR products were pooled, purified using a MinElute PCR Purification Kit (QIAGEN, Germany), and electrotransformed into Endura electrocompetent cells (Biosearch Technologies, USA). To ensure comprehensive coverage, parallel transformations were conducted, and colony numbers were recorded to achieve 1,000-fold coverage of the total number of sgRNAs in the library. Subsequently, the sgRNA plasmid library was extracted using a Plasmid Plus Maxi Kit (QIAGEN, Germany). The sgRNA cassettes from the plasmid library were amplified with PrimeSTAR GXL DNA Polymerase (Takara, China) as previously described. The PCR products were purified using a MinElute PCR Purification Kit (QIAGEN, Germany) and sequenced on an Illumina HiSeq2000 platform. The coverage and homogeneity of sgRNAs were analyzed using the MAGeCK algorithm. All primers used for constructing the sgRNA expression vector are listed in Supplementary Table 4.

### 4.7 Lentivirus production

HEK293T cells were seeded in T25 flasks (Thermo Fisher Scientific, 156367) at approximately 30% confluency one day prior to transfection. On the following day, when the cells reached 90–99% confluency, transfection was performed. For each T25 flask, the transfection mixture consisted of 2.4 µg of the plasmid containing the vector of interest, 2.4 µg of psPAX2 (Addgene, 12260), and 1.2 µg of pMD2.G (Addgene, 12259). This mixture was transfected using 18 µL of Lipofectamine® LTX Reagent (Invitrogen, Thermo Fisher Scientific, USA, 15338030), 6 µL of Lipofectamine® LTX PLUS Reagent (Thermo Fisher Scientific, USA, 15338100), and 1.25 mL of Opti-MEM (Thermo Fisher Scientific, USA, 31985070). Transfection parameters were scaled linearly with flask surface area for T75 and T225 flasks. The culture medium was replaced 6 hours post-transfection. The virus-containing supernatant was harvested 48 hours later, filtered through a 0.45-µm PVDF filter (Millipore Sigma, SLHV013SL), and concentrated by ultracentrifugation at 100,000 × g for 2 hours at 4 ℃ when necessary. The virus supernatant was then aliquoted and stored at −80 ℃.

### 4.8 ORFV infection of the ovine genome-wide CRISPR/Cas9 knockout cell library and flow cytometry sorting

To generate ovine genome-wide CRISPR/Cas9 knockout cell libraries, a total of 4 × 10^8^ OA3.Ts/Cas9 cells were transduced with lentivirus of ovine genome-wide CRISPR/Cas9 knockout sgRNA library in the presence of 0.6 mg/ml polybrene (107689, Sigma), at a multiplicity of infection (MOI) of 0.2. Two days post-transduction, transduced cells were selected with 1 μg/ml puromycin (Coolaber, CP9231) for 10-14 days, while maintaining at least a coverage of 1000x. Subsequently, a library of 1.2×10^8^ OA3.Ts/Cas9 cells was seeded into 225 cm² cell culture dishes (Thermo Scientific, 130330) and infected with ORFV at a multiplicity of infection (MOI) of 2. At 48 hours post-infection (hpi), cells were collected, fixed and permeabilized using the reagent from Cytofix/Cytoperm™ Fixation/Permeabilisation Kit(BD, 554714). After washing three times with Perm/Wash™ Buffer(BD, 554723), cells were incubated with ORFV antibody(Antibodies-online, ABIN6774766) for 1 hour, followed by another three washes with Wash Buffer. Cells were then incubated with Alexa Fluor 488-labelled Goat Anti-Mouse IgG(H+L) (Beyotime, A0428) for 30 minutes and washed three times with Wash Buffer. Finally, cells were resuspended in PBS containing 1% BSA and subjected to cell sorting to collect the top 5% of GFP-expressing cells. Flow cytometry was performed using a Sony MA900 FACS Flow Cytometry System and MA900 Cell Sorter software.

### 4.9 Screen analysis

Read counts for each guide RNA were normalized to reads per million and log-transformed. Quantile normalization was performed in R. To account for the significant heteroscedasticity (Figure 1b), local z-scores were calculated for groups of values with varying read counts, using sliding bins of different sizes. For any comparison of two samples from which n read counts[x] and [y] are derived (for example, the ORFV-permissive and control FACS pools). Z-scores were then calculated within each of these bins. *P*-values were calculated from the sum of z-scores for sgRNAs targeting a particular gene compared to a density function modelled on an empirical distribution of possible combinations of sgRNA z-scores permuted at least 1e8 times by randomly rearranging z-scores for all sgRNAs in the screen. In order to minimise false negatives and maximise the discovery power of our screen, we did not require more than one sgRNA per gene to be significantly over-represented in the ORFV-permissive FACS pool (permissive set). We report an additional “robust” set of hits in which the empirical p-value for a given gene, derived from the remaining sgRNAs after the sgRNA with the most significant effect is removed (remainder p), is less than 0.05. FDRs were calculated using the Benjamini-Hochberg method in scipy stats v1.1.0.

### 4.10 Cell viability assay

The cells were seeded into opaque-walled 96-well plates at a density of 5 × 10^3^ cells per well in 100 µl of DMEM and incubated for 24 h. The CCK-8 assay (Dojindo, Kumamoto, Japan) was performed to analyze cell viability at 0, 24, 48, 72, and 96 h. The cells were incubated with the CCK-8 reaction solution for two hours at 37 °C. A SpectraMax Mini microplate reader (Molecular Devices, USA) was employed to measure the absorbance at 450 nm, and the optical density (OD) values were obtained to assess the cells’ proliferation abilities.

### 4.11 Generation of PEX11B knockout OA3.Ts/Cas9 single-cell clone

The construction of the lentivirus sgRNA expression vector was initiated by digesting the lenti-pcg2.0 vector with the BbsI restriction enzyme. The paired oligonucleotides of sgRNA were annealed and subsequently cloned into the linearised vector. The oligonucleotides (PEX11B oligo F: ACCGACAGAAACAGATTCGACAAC and PEX11B oligo R: AAACGTTGTCGAATCTGTTTCTGT) were synthesized (Beijing Genomics Institution) and annealed in a 50 μl reaction containing TransTaq HiFi Buffer II at a final concentration of 9 μM. The annealed oligonucleotides were ligated into a lentiviral vector carrying a puromycin selection marker using Golden Gate Assembly (NEB). The ligation products were then transformed into Trans1-T1 competent cells (Transgen, CD501). Pseudotyped lentiviral vectors expressing sgRNA were generated by following the previously described lentivirus production steps. The OA3.Ts/Cas9 cell line was transduced with these pseudotyped lentiviral vectors and selected with puromycin (1 mg/ml). Single-cell clones resistant to puromycin were selected. Genomic DNA was extracted from these clones, and the insertions and deletions (indels) caused by sgRNA/Cas9 in each cell clone were confirmed by Sanger sequencing after PCR amplification. The monoclonal cells that stably expressed the Cas9 gene and exhibited the highest knockout efficiency were obtained through Western blot and knockout efficiency detection (TIDE: http://tide.nki.nl/).

### 4.12 Generation of PEX11B overexpression and mutant OA3.Ts/Cas9 cell line

To construct the overexpression vector for the generation of PEX11B stable cell lines, the coding sequences of PEX11B were cloned into the pLV3-CMV-MCS-3×Myc-Neo vector (Clontech), which was linearised with BamHI. Subsequently, to abolish cleavage by sgRNA and Cas9 in KO-PEX11B cells, a specific point mutation in protospacer adjacent motif sequences, which does not alter the amino acid, was introduced into the PEX11B coding sequences to construct its mutant overexpression vectors. The PEX11B mutant was generated following the same procedure as described above. Sanger sequencing (Tsingke) was performed to confirm all plasmids. All primer sequences are listed in Supplementary Table 4.

### 4.13 RNA isolation and RT-qPCR

The cells were lysed in TRIzol (Tiangen), and RNA was isolated according to the manufacturer’s protocol. The cDNA was synthesized using the Evo M-MLV RT Premix for qPCR (Accurate Biology, 11728). RT-qPCR was performed using TB Green® Premix Ex Taq™ II FAST qPCR (Takara, CN830A). Gene expression was quantified using the 7500 Real-Time PCR System (Thermo Fisher Scientific). The sequences of the primers targeting ORFV B2L were as follows: GGGCTCTACTCCACCAACAA (forward) and CGAGTCCGAGAAGAATACGC (reverse). The expression of target genes was normalisednormalized to β-actin.were as follows: CCATCGTCCACCGCAAAT (forward) and CAAATAAAGCCATGCCAATCTC (reverse).

### 4.14 ORFV infection and purification

OA3.Ts/Cas9 cells were inoculated with 250 μL (6-well plate) or 2 mL (T75 flask) of ORFV at MOI 2 for 1.5 hours at 37℃ in serum-free DMEM. The cells were then washed and replaced with fresh serum-free DMEM supplemented for 48 h. The TCID50 assay subsequently monitored infection. The ORFV was concentrated using the Universal Virus Concentration Kit (Beyotime #C2901S), following the manufacturer’s instructions. To summarise, the viral supernatant was initially collected, and cellular debris was removed through centrifugation and filtration. The pre-cooled Virus Precipitation Reagent was then mixed in proportion with the viral supernatant, thoroughly mixed, and stirred at a low speed at the appropriate temperature overnight. This was followed by centrifugation to remove the supernatant and careful aspiration of the precipitate. The process was then repeated, and the supernatant was collected as concentrated virus.

### 4.15 R18 Loading of ORFV Particles

100 µl of purified ORFV was added to 2 µl of 5 µmol Octadecyl Rhodamine B Chloride (R18; MKBio, 65603-19-2) in phosphate-buffered saline (PBS; Quality Biological) + 0.2% bovine serum albumin (Thermo; 10010023) for 60 min at room temperature in the dark. Non-incorporated R18 was removed by a 0.45 μm sterile filter (MerckMillipore, SLHVR33RB).

### 4.16 R18-ORFV binding assay

Cells were seeded on a 40mm confocal dish(Thermofisher, 150680) and inoculated with R18-ORFV at an MOI of 100 for 60 minutes at 4 °C. Cells were then washed twice with ice-cold PBS to remove unbound virus and fixed with 4 % paraformaldehyde(Beyotime, P0099). Following a 10-minute incubation with hochest 33258 (Thermo, H1398)staining solution at room temperature, the cells were washed three times with PBS and subsequently imaged with a confocal microscope.

### 4.17 Measuring lysosomal degradation of DQ-BSA

1×10^5^ OA3.Ts/Cas9 cells were seeded on a 40mm confocal dish (Thermofisher, 150680) and incubated with 10ug/ml DQ Green BSA (Sharebio, D-12050SB). Following 8 hours of standard culture, the dye-containing culture medium was subsequently replaced. 200 μL DAPI for one hour at 37℃. The cells were then washed in PBS and fixed in 4% paraformaldehyde. Confocal microscopy Images(Zeiss LSM 710 Confocal Microscope) were acquired with a ×40 objective.

### 4.18 TCID50 assay

OA3.Ts/Cas9 cells were seeded at a density of 1×10⁵ cells per well in 96-well plates and cultured for a period of 24 hours. The ORFV to be tested was thawed on ice and diluted at a 10-fold multiplicity of infection, with eight wells per dilution. The medium in the cell plates was aspirated, washed three times with PBS, and incubated with 100 μL of virus solution for one hour. The venom was aspirated, washed three times with PBS, and 200 μL of 2% serum-containing DMEM medium was added. The cell lesions were observed on the fifth day after infection.

### 4.19 LC-MS

The mass spectrometry signal acquisition of the sample was performed in separate positive and negative ion modes. The specific acquisition mode employed was data-dependent acquisition (DDA). Non-target cells and KO-PEX11B Cells were transferred into a 2 mL centrifuge tube with 600 μL methanol-water (V: V=1:1). An internal standard (Lyso PC-17:0, 0.1 mg/mL, prepared in methanol) was added at a volume of 20 μL. Subsequently, 600 μL chloroform was added to the mixture. The sample was subjected to sonication in an ice bath at 500 W for 3 min with a cycle of 6 s on and 4 s off. After that, the sample was subjected to ultrasonic extraction in an ice-water bath for 10 min. The mixture was then left to stand at 4℃ for 30 min and centrifuged at 12000 rpm for 10 min at 4℃. The lower phase (400 μL) was collected and transferred to an LC-MS vial for evaporation. In the remaining centrifuge tube, 600 μL chloroform-methanol (V: V=2:1) was added. The mixture was vortexed for 30 s and subjected to ultrasonic extraction in an ice-water bath for 10 min. The mixture was left to stand at 4℃ for 30 min and then centrifuged at 12000 rpm for 10 min at 4℃. The lower phase (400 μL) was collected and transferred to the same LC-MS vial for further evaporation. The lipid residue in the LC-MS vial was re-dissolved in 300 μL isopropanol-methanol (V: V=1:1) by vortexing for 30 s and sonicating in an ice-water bath for 3 min. The solution was then transferred to a 1.5 mL EP tube and centrifuged at 12000 rpm for 10 min at 4℃. The supernatant (200 μL) was transferred to an LC-MS vial with an insert for LC-MS analysis of Ultra-High Performance Liquid Chromatography Tandem High-Resolution Mass Spectrometer (Dionex U3000 UHPLC; UHPLC-HRMS/MS). The quality control (QC) sample was prepared by mixing equal volumes of the extraction solutions from all samples, with each QC sample having the same volume as the individual samples. Note: All extraction solvents were pre-cooled at −20℃ before use.

### 4.20 Fluorescence recovery after photobleaching (FRAP) assay

For FRAP experiments, cells have been seeded in a 40mm confocal dish (Thermofisher, 150680) with 2 × 10^5^ cells per dish the day before the experiment. The solution has been removed completely, and 100 μL of DiOC18 (MKBio, MX4001)in phosphate-buffered solution has been added. After 15 min of incubation at 37°C, cells were washed three times with PBS. FRAP measurements were performed on a ZISSE LSM710 confocal microscope using a 40x water objective. The incubation chamber around the microscope was set to 37°C. Imaging was performed with the 488-nm laser line of an argon laser set to 70% laser power with less than 10% transmission. Images were acquired at a rate of 1.318 s/image. Five frames were recorded before bleaching and 72-100 after bleaching. Bleaching was performed by zooming in on the bleaching area and scanning once with all lines of the argon laser at 100% transmission, plus a UV diode at 100% transmission. This resulted in bleaching of > 80% of the fluorescence intensity in the bleached spot. For analysis, the background fluorescence outside of the cells was first subtracted from the image. The intensity in the bleached spot was normalized to the intensity in the whole cell to correct for the total loss of fluorescence during the experiment. It was further normalized to pre-bleaching intensity in the spot.

The mobile fraction (*mf*) was calculated using the asymptote (*y_0_*) of this function:

*mf* =*(y0* - *Ibleached / Iprebleached-Ibleached)*

Where Ibleached is the fluorescence intensity in the bleached area in the first frame after bleaching, and Iprebleached is the mean intensity in that area in the five frames before bleaching.Accordingly, the immobile fraction (*If*) is:

*If=1-mf*

The diffusion constant (*D*) was calculated using the time to reach half maximum (τ_1/2_) of the exponential fit and the equation for two-dimensional diffusion:

*D*=R^2^/4τ_1/2_

### 4.21 Fluorescence anisotropy evaluation

The fluorescence anisotropy of 1-[4-(trimethylamino)phenyl]-6-phenyl-1,3,5-hexatriene (TMA-DPH, MCE, HY-D0986) incorporated in the cells was assessed by the determination of steady state fluorescence polarisation of the membrane-fluorescent probe system; the TMA-DPH probe lacks fluorescence in solution and becomes fluorescent when incorporated into the lipid membrane bilayer. 1 × 10^5^ cells were collected and suspended in a solution containing one µM TMA-DPH. The suspension was incubated at 37℃ for 20 minutes, followed by centrifugation at 800 rpm for 3 minutes. The supernatant was discarded, and the cells were resuspended in 2 mL of PBS (pH 7.4). The cell suspension was then aliquoted at 150 µL per well into a black microplate for detection with a Spectra Max iD5 Microplate detection instrument with blue polarised light detection (excitation wavelength 355 nm; emission maximum 430 nm) at room temperature. Calculation of the fluorescence anisotropy (R) was performed according to equations (A) and (B):

A. R=(*I*_VV_*-GI*_V0_)/(*I*_VV+_*GI*_V0_)
B. G=*I*_0V_/*I*_00_

Where R is the fluorescence anisotropy, I_vv_, I_vo_, I_ov_ and I_oo_ represent the emission intensity corrected for the autofluorescence signal of unstained cells, when the polarisers in the excitation end emission beams are oriented in vertical-vertical, vertical-horizontal, horizontal-vertical and horizontal-horizontal positions, respectively.

### 4.22 Confocal microscopy

For 100ul R18-ORFV with 900 µl of PBS, and incubated under light-avoiding room temperature conditions for one h. At the conclusion of the labelling process, any unbound R18 was filtered out using a 0.45 μm filter. Cells were subjected to three washes with PBS, after which the labelled virus was added and allowed to bind at 4 ℃ for one hour. At the conclusion of the procedure, any unbound virus was removed by washing with PBS, and the cells were incubated at 37 ℃ for the indicated times to allow for internalization (early endosomes: 10 minutes, late endosomes: 20 minutes, lysosomes: 30 minutes). The cells were fixed using 4% paraformaldehyde for a period of 10 minutes. Subsequently, the cells were punched using Immunostaining Permeabilisation Buffer with Saponin (Beyotime#P0095), in accordance with the instructions provided by the manufacturer. The second antibodies were used in a 1:300 configuration: anti-Rab5 (Immunoway#YT5456), Rab7 (Abcam#ab126712), and Lamp1 (Proteintech#65051). Cells were incubated overnight. Following the application of the anti-rabbit secondary antibody (Proteintech#SA00003) in a 1:200 configuration, the cells were incubated for a period of 2 hours. Subsequently, the cells were stained with Hoechst 33258 Staining Solution for 10 minutes. The images were captured using a Zeiss LSM 710.

### 4.23 3D reconstructions

All peroxisome-related microscopy work was performed using a Nikon AXR confocal microscope system. This integrated system incorporates a high-speed resonant scanning confocal unit, a high-sensitivity sCMOS camera, and a high-precision motorised stage optimised for advanced imaging applications. Images (1024×1024 pixels) were captured on a Nikon AXR confocal microscope using a 100×oil objective with a 1.4 numerical aperture. Z-steps of 0.2 microns were used for analysis of peroxisome morphology and colocalization. FIJI (Fiji Is Just ImageJ)software was used to classify surfaces, eliminate noise in each channel, and create 3D reconstructions.

### 4.24 Gene editing efficiency testing by T7E1 digestion and TIDE analysis

Seven to eight days after transfection with lentivirus or individual CRISPR sgRNAs, crude genomic DNA was extracted from cells using QuickExtract DNA Extraction Solution following the product manual. The extract was diluted 1:10 in nuclease-free water, and one μL of the dilution was used for PCR using primers flanking the target site and Phusion HF polymerase with supplied buffer, producing amplicons between 300- 800bp in size. 2 μL of the PCR reaction was run on 2% agarose gel to estimate the amplicon concentration. Without purification, the calculated volume of PCR reaction containing ∼200ng PCR product based on the agarose gel was denatured at 95℃ for 5 minutes and re-annealed by dropping the temperature to 25℃at 0.1℃ per second in a thermal cycler. At the end of the program, the temperature was reduced to 4℃. Right after re-annealing, one μL T7 Endonuclease I was added to the PCR reaction directly and incubated at 37℃ for 30 minutes. Immediately after incubation, the reaction was analysed on a 2% agarose gel. The Densitometry function of ImageJ processed the gel image, and the percentage editing was calculated. The editing efficiency was also examined by TIDE analysis (https://tide.deskgen.com/) following Sanger sequencing of purified PCR products.

### 4.25 Detection of IMV (immature virions) and MV (mature virions) during ORFV infection using transmission electron microscopy

The process of ORFV infection was observed using transmission electron microscopy. Briefly, non-target cells and KO-PEX11B cells were infected with viruses (MOI 1) and incubated at 4 °C for 1 h. After incubation, PBS was used to wash away unbound viruses. The bound virus was then allowed to infect at 37 °C for 48 hours. Following infection, the cells were fixed with 2.5% glutaraldehyde for 16 h. Finally, ultrathin sections were prepared on carbon-coated 100-mesh copper grids and observed in a Hitachi-7650 transmission electron microscope at an operating voltage of 80 kV.

### 4.26 Statistical analysis

Statistical analyses were conducted using GRAPHPAD Prism 9.0 software(La Jolla, CA, USA). All experiments were repeated at least three times, and the distribution of data points is presented as mean SEM. Results are shown as means ± standard deviations (SD) of the data obtained from three independent experiments or triplicates as indicated. Comparisons between samples were done using the paired two-tailed t-test. *P* < 0.05 was considered significant.

## Acknowledgements

This project was supported by a grant from the Identifying Key Host Factors for Orf Virus Infection via Sheep CRISPR Whole-Genome Screen and Elucidating the Antiviral Mechanism of PEX11B (32560851),the Inner Mongolia Autonomous Region Major Science and Technology Project (2021ZD0013), the ‘Grassland Talents’ Program of Inner Mongolia Autonomous Region (12000-12102617), the High-Level Talents Research Support Program of Inner Mongolia Autonomous Region (10000-21311201/004), the ‘Horse Program’ High-Level Talents Program of Inner Mongolia University (10000-21311201/141), the National Key Laboratory of Reproductive Regulation and Breeding of Grassland Livestock (Jointly Built by the Province and Ministry) - Identification of Specific Target Points for Important Pathogens in Cattle and Sheep and Development of Novel Diagnostic Technologies(2025KYPT0066), the Science and Technology Leading Talent Team Program of Inner Mongolia Autonomous Region (2022LJRC0009 to W. Hu from Inner Mongolia University and Fudan University), the Science and Technology Major Special Project of Inner Mongolia Autonomous Region (2020ZD0008 to Prof. Wei Hu from Inner Mongolia University and Fudan University).

## Author contributions

G.J.W. and W.H. conceived and designed the experiments; X.R.G.,Y.J.H.,S.L.,J.F.W.,Y.S., X.K. X.G., Y.S.,Y.Z.S., Y.T and Y,W.Y. performed the experiments; X.R.G., and Y.J.H.analyzed the data; Q.Y.Z. provided the viruses. J.F.W.,and S.D.W carried out electron-microscopy imaging.G.J.W.,and R.H., J.L.W. interpreted the data; X.R.G. and G.J.W. wrote the original draft; G.J.W., and W.H. reviewed and edited the paper.

## Supplemental information

### Supplemental figures

Figure S1. Validation of the successful construction of pLV-cas9-hTERT plasmid (Related to Figure 1). (A) The pLV-cas9-hTERT plasmid after double digestion with BsmBI and EcoRI.(1: pLV-cas9-hTERT plasmid digested with BsmBI enzyme; 2: pLV-cas9-hTERT plasmid digested with EcoRI enzyme; 3: pLV-cas9-hTERT plasmid digested with both BsmBI and EcoRI enzymes; 4: undigested pLV-cas9-hTERT plasmid.) (B) pLV-cas9-hTERT lasmid sequencing diagram, showing the sequencing of the recombinant connection part.

Figure S2. Verification of immortalized OA3.Ts/Cas9 cells (Related to Figure 1). (A) Determination of lentiviral infection efficiency in OA3.Ts/Primary cells. (B) Identification of Cas9 and hTERT genes in OA3.Ts/Cas9 cells. (C) Morphology of OA3.Ts/Cas9/P107 cells. (D) Cell proliferation detection at different passage numbers. (E) (a)(b)Detection of apoptosis efficiency in cells of different passage generations.

Figure S3. Establishment of a monoclonal Cas9-expressing line by single-cell cloning (Related to Figure 1). (A) Cas9 gene expression in OA3.Ts/Cas9 detected by qPCR. (B) hTERT gene expression in OA3.Ts/Cas9 detected by qPCR. (C) Evaluation of Cas9 activity among candidate single–cell–derived clones using a T7EN I cleavage assay. The candidate cells were transduced with a validated sgRNA (targeting the B4GALNT2 gene) lentivirus. The single–cell–derived clone with the highest Cas9 activity is Clone#A. (D) (a-h)The TIDE tool quickly determines the type and proportion of mutations in gene editing by comparing the Sanger sequencing peak profiles of PCR products before and after editing.

Figure S4. Evaluation of the cleavage efficiency of Cas9 in OA3.Ts/cas9/clone#A cells (Related to Figure 1). (A) Cas9 and hTERT expression does not affect ORFV infection in OA3.Ts/cas9 clone. (B) Assessment of the transduction efficiency of sgRNA lentivirus in OA3.Ts/cas9/clone#A cells at the time points indicated. (C) Assessment of the cleavage activity of sgRNA lentivirus in OA3.Ts/cas9/clone#A cells at the time points indicated using a T7EN I assay. Indel%: percentage of indels; bp: base pairs; dpi: days post infection, Control: wild-type cells; Marker: Marker I DNA ladder. Scale bar, 200 μm. (D) Assessment of gene editing efficiency in OA3.Ts/Cas9/clone#A cells using the T7E1 endonuclease digestion assay. Results indicate that the gene editing efficiency tends to stabilize at 6 days following lentiviral infection.

Figure S5. Generation of ovine genome-wide sgRNA cell libraries (Related to Figure 2). (A) Overview table of distribution of ovine genome-wide sgRNA Library. (B) The count of mismatches for each sgRNA in the library. The information is derived from the CRISPR sgRNA library of the sheep whole genome, which is utilized to evaluate the targeting precision and off-target potential of sgRNAs. (C) Sequencing result of sgRNAs sequence distribution accumulation in CRISPR pooled sgRNA plasmid pools. (D) Evaluation of knockout effects of a randomly selected sgRNA from the initially designed sgRNA library. (E) Analysis of lentiviral infection titer in the genome-wide library. (a) Analysis of the positive rate of cells following lentiviral infection (with 5-fold serial dilution) using flow cytometry.(b) Analysis of the positive rate of cells following lentiviral infection (with 5-fold serial dilution) using fluorescence microscopy.

Figure S6. Identification of the viral potential enhanced in clonal RAD52, USP45, CDH13, LRP12, PEX11B, GJB6, and SLC12A5(Related to Figure 3). (A) (a,b)Genomic editing efficiency of selected CRISPR sgRNAs measured by massively parallel sequencing of the CRISPR target site and TIDE analysis. (B) The CPE (cytopathic effect) of seven-knockout cell lines infected at different time points.(MOI=2)

Figure S7. Validation of PEX11B as a host restriction factor against ORFV infection(Related to Figure 4). (A) Cell viability in KO-PEX11B versus non-target cells by cell counting kit-8 assay. (B) Relative quantitative real-time PCR detection of PEX11B expression levels. (C) Bioinformatics assessment of potential off-target genomic sites using CRISPR/Cas9 and our sgRNA targeting PEX11B: PCR-amplification followed by sequencing of genomic DNA extracted from KO-PEX11B cells confirmed that these potential off-target sites showed wild-type sequences. The cleavage efficiency of T7EN I at this potential off-target site is shown in (D). (D) Off-target cleavage efficiencies at seven predicted loci were quantitatively assessed by T7E1 nuclease digestion of corresponding PCR amplicons. (E) Confocal microscopy analysis of ORFV in Non-target versus KO-PEX11B cell lines 90 min after infection. Blue, Hoechst 33258; red, R18-labeled ORFV; cell n=45. (F) Analysis of endocytic fusion of ORFV infection. After incubating ORFV at a multiplicity of infection (MOI=1) with the cell line at 4 ℃ for 2 hours, the cells were washed with ice-cold PBS at pH 7.4 and then incubated at 37 ℃ for 1 hour. Subsequently, uninternalized viruses were washed away with PBS at pH 3.0. After incubating at 37 degrees Celsius for 24 hours, immunofluorescence assays were performed.

Figure S8. Construction and Validation of PEX11B Mutant Cell Lines(Related to Figure 5). (A) Cell viability of non-target cells treated with GW6471 was assessed using the Cell Counting Kit-8. (B) Cell viability of non-target cells treated with Wy14643 was assessed using the Cell Counting Kit-8.

Figure S9. PEX11B ablation elicits peroxisome remodeling in morphology and size. (Related to Figure 6). (A) Retrovirus-induced gene complement of Myc-tagged PEX11B mutants (△159-183,△185-210,△210-259) in the KO-PEX11B clonal cell line. (B) Quantitative analysis of ORFV infection. Flow cytometry analysis was performed on cell lines infected with ORFV (MOI=1) using ORF 086 antibodies. (C) Maximum projections of non-target and KO-PEX11B cells with anti-PMP70 and imaged at 100X. A square frame indicates ROIs in the corresponding color. Scale bars=10µm. (D) Plot of the surface area and volume of individual peroxisomes in (a) Non-target cells (mean SA: V = 23.89 μm-1), (b) KO-PEX11B (mean SA: V = 27.64 μm-1). Solid line indicates the regression curve, dashed lines indicate the upper and lower 95% confidence intervals. N = 23868 peroxisomes in Non-Target; N = 7391 in KO-PEX11B.

Figure S10. PEX11B deletion reprograms the lipidomic landscape of viral and cellular membranes. (A) Statistical summary of lipidomics categorization of lipids in cells. Lipid relative abundances were quantified based on peak area normalization. The resultant lipidome comprised six major classes: fatty acyls (FA), glycerolipids (GL), glycerophospholipids (GP), sphingolipids (SL and SP), and a minor heterogeneous lipid fraction designated as “Class.” (B) Principal component analysis (PCA) score plot of lipidomic profiles. (C) Permutation test (n = 200) validating the OPLS-DA model. The plot displays R² (green triangle) and Q² (blue square) values obtained after 200 random permutations of the Y matrix. The original model’s R² = 0.997 and Q² = −0.055 (red symbols) fall outside the permutation distribution, indicating that the observed separation is not due to over-fitting and the model possesses robust predictive power. (D) Differential lipid distribution volcano plot. Red dots represent metabolites that are upregulated in the experimental group, blue dots represent downregulated metabolites, and gray dots represent metabolites that are not significant. The horizontal axis shows the log2(FC) values for the comparison between the two groups, while the vertical axis represents -log10(p-value) values. Individual significant differences in lipids have been annotated. (F)A classified bubble chart showing the relationship between carbon chain length and unsaturation in PC(a), PE (b), and SM(c). The x-axis represents the carbon chain length of the lipids, while the y-axis represents the unsaturation of the lipids. The color of the bubbles maps to Log2(FC), and larger circles indicate smaller *P*-value values.

### Supplemental tables

Table S1. MAGeCK Analysis of CRISPR Screens. Related to **Figure 2**.

This supplementary table (Table 1) presents the sgRNA sequences utilized for the construction of the ovine genome-wide CRISPR library.

Table S2. MAGeCK Analysis of CRISPR Screens. Related to Figure 3.

This supplementary table (Table 2) presents the sequencing results for sgRNAs targeting sequences within CRISPR-knockout, sorted mutant cell populations that contain the complete sgRNA library.

Table S3. Gene ontology analysis of 0.1% of the ranked hits from the result of the MAGeCK analysis. Related to Figure 3.

This supplementary table (Table 3) presents the Gene Ontology (GO) analysis of 0.1% of the ranked hits from the MAGeCK analysis results, highlighting enriched biological processes, molecular functions, or cellular components associated with the identified host genes in the context of ORFV screening.

Table S4. Primers used in this research and related to STAR Methods.

This supplementary table (Table 4) lists the primers used in this research, corresponding to the details described in the Methods.

## References

1. Bergqvist C, Kurban M, Abbas O. Orf virus infection. Rev Med Virol. 2017;27(4). Epub 20170508. doi: 10.1002/rmv.1932. PubMed PMID: 28480985.

2. Wang R, Wang Y, Liu F, Luo S. Orf virus: A promising new therapeutic agent. Rev Med Virol. 2019;29(1):e2013. Epub 20181028. doi: 10.1002/rmv.2013. PubMed PMID: 30370570.

3. Spyrou V, Valiakos G. Orf virus infection in sheep or goats. Vet Microbiol. 2015;181(1-2):178–82. Epub 20150812. doi: 10.1016/j.vetmic.2015.08.010. PubMed PMID: 26315771.

4. da Costa RA, Cargnelutti JF, Schild CO, Flores EF, Riet-Correa F, Giannitti F. Outbreak of contagious ecthyma caused by Orf virus (Parapoxvirus ovis) in a vaccinated sheep flock in Uruguay. Braz J Microbiol. 2019;50(2):565–9. Epub 20190305. doi: 10.1007/s42770-019-00057-7. PubMed PMID: 30835059; PubMed Central PMCID: PMCPMC6863263.

5. Delhon G, Tulman ER, Afonso CL, Lu Z, de la Concha-Bermejillo A, Lehmkuhl HD, et al. Genomes of the parapoxviruses ORF virus and bovine papular stomatitis virus. J Virol. 2004;78(1):168–77. doi: 10.1128/jvi.78.1.168-177.2004. PubMed PMID: 14671098; PubMed Central PMCID: PMCPMC303426.

6. Rintoul JL, Lemay CG, Tai LH, Stanford MM, Falls TJ, de Souza CT, et al. ORFV: a novel oncolytic and immune stimulating parapoxvirus therapeutic. Mol Ther. 2012;20(6):1148–57. Epub 20120124. doi: 10.1038/mt.2011.301. PubMed PMID: 22273579; PubMed Central PMCID: PMCPMC3369287.

7. Abu Ghazaleh R, Al-Sawalhe M, Abu Odeh I, El Ibrahim J, Al-Turman B, Makhamreh J. Host range, severity and trans boundary transmission of Orf virus (ORFV). Infect Genet Evol. 2023;112:105448. Epub 20230520. doi: 10.1016/j.meegid.2023.105448. PubMed PMID: 37217030.

8. van Vloten JP, Matuszewska K, Minow MAA, Minott JA, Santry LA, Pereira M, et al. Oncolytic Orf virus licenses NK cells via cDC1 to activate innate and adaptive antitumor mechanisms and extends survival in a murine model of late-stage ovarian cancer. J Immunother Cancer. 2022;10(3). doi: 10.1136/jitc-2021-004335. PubMed PMID: 35296558; PubMed Central PMCID: PMCPMC8928368.

9. Bukar AM, Jesse FFA, Abdullah CAC, Noordin MM, Lawan Z, Mangga HK, et al. Immunomodulatory Strategies for Parapoxvirus: Current Status and Future Approaches for the Development of Vaccines against Orf Virus Infection. Vaccines (Basel). 2021;9(11). Epub 20211117. doi: 10.3390/vaccines9111341. PubMed PMID: 34835272; PubMed Central PMCID: PMCPMC8624149.

10. Hosamani M, Scagliarini A, Bhanuprakash V, McInnes CJ, Singh RK. Orf: an update on current research and future perspectives. Expert Rev Anti Infect Ther. 2009;7(7):879–93. doi: 10.1586/eri.09.64. PubMed PMID: 19735227.

11. Sevik M. Orf virus circulation in cattle in Turkey. Comp Immunol Microbiol Infect Dis. 2019;65:1–6. Epub 20190326. doi: 10.1016/j.cimid.2019.03.013. PubMed PMID: 31300096.

12. Haig DM, Mercer AA. Ovine diseases. Orf. Vet Res. 1998;29(3-4):311–26. PubMed PMID: 9689744.

13. Nagendraprabhu P, Khatiwada S, Chaulagain S, Delhon G, Rock DL. A parapoxviral virion protein targets the retinoblastoma protein to inhibit NF-kappaB signaling. PLoS Pathog. 2017;13(12):e1006779. Epub 20171215. doi: 10.1371/journal.ppat.1006779. PubMed PMID: 29244863; PubMed Central PMCID: PMCPMC5747488.

14. Zhao K, Song D, He W, Lu H, Zhang B, Li C, et al. Identification and phylogenetic analysis of an Orf virus isolated from an outbreak in sheep in the Jilin province of China. Vet Microbiol. 2010;142(3-4):408–15. Epub 20091031. doi: 10.1016/j.vetmic.2009.10.006. PubMed PMID: 19948384.

15. Imlach W, McCaughan CA, Mercer AA, Haig D, Fleming SB. Orf virus-encoded interleukin-10 stimulates the proliferation of murine mast cells and inhibits cytokine synthesis in murine peritoneal macrophages. J Gen Virol. 2002;83(Pt 5):1049–58. doi: 10.1099/0022-1317-83-5-1049. PubMed PMID: 11961259.

16. Li S, Jing T, Zhu F, Chen Y, Yao X, Tang X, et al. Genetic Analysis of Orf Virus (ORFV) Strains Isolated from Goats in China: Insights into Epidemiological Characteristics and Evolutionary Patterns. Virus Res. 2023;334:199160. Epub 20230705. doi: 10.1016/j.virusres.2023.199160. PubMed PMID: 37402415; PubMed Central PMCID: PMCPMC10410590.

17. Tang X, Xie Y, Li G, Niyazbekova Z, Li S, Chang J, et al. ORFV entry into host cells via clathrin-mediated endocytosis and macropinocytosis. Vet Microbiol. 2023;284:109831. Epub 20230713. doi: 10.1016/j.vetmic.2023.109831. PubMed PMID: 37480660.

18. Khatiwada S, Delhon G, Chaulagain S, Rock DL. The novel ORFV protein ORFV113 activates LPA-p38 signaling. PLoS Pathog. 2021;17(10):e1009971. Epub 20211006. doi: 10.1371/journal.ppat.1009971. PubMed PMID: 34614034; PubMed Central PMCID: PMCPMC8523077.

19. Li Q, Brass AL, Ng A, Hu Z, Xavier RJ, Liang TJ, et al. A genome-wide genetic screen for host factors required for hepatitis C virus propagation. Proc Natl Acad Sci U S A. 2009;106(38):16410–5. Epub 20090827. doi: 10.1073/pnas.0907439106. PubMed PMID: 19717417; PubMed Central PMCID: PMCPMC2752535.

20. Ayres JS, Schneider DS. Tolerance of infections. Annu Rev Immunol. 2012;30:271–94. Epub 20120103. doi: 10.1146/annurev-immunol-020711-075030. PubMed PMID: 22224770.

21. Cornel MC, van der Meij KRM, van El CG, Rigter T, Henneman L. Genetic Screening-Emerging Issues. Genes (Basel). 2024;15(5). Epub 20240503. doi: 10.3390/genes15050581. PubMed PMID: 38790210; PubMed Central PMCID: PMCPMC11121342.

22. Kool J, Berns A. High-throughput insertional mutagenesis screens in mice to identify oncogenic networks. Nat Rev Cancer. 2009;9(6):389–99. doi: 10.1038/nrc2647. PubMed PMID: 19461666.

23. Wei J, Alfajaro MM, DeWeirdt PC, Hanna RE, Lu-Culligan WJ, Cai WL, et al. Genome-wide CRISPR Screens Reveal Host Factors Critical for SARS-CoV-2 Infection. Cell. 2021;184(1):76–91 e13. Epub 20201020. doi: 10.1016/j.cell.2020.10.028. PubMed PMID: 33147444; PubMed Central PMCID: PMCPMC7574718.

24. Flint M, Chatterjee P, Lin DL, McMullan LK, Shrivastava-Ranjan P, Bergeron E, et al. A genome-wide CRISPR screen identifies N-acetylglucosamine-1-phosphate transferase as a potential antiviral target for Ebola virus. Nat Commun. 2019;10(1):285. Epub 20190117. doi: 10.1038/s41467-018-08135-4. PubMed PMID: 30655525; PubMed Central PMCID: PMCPMC6336797.

25. Ma H, Dang Y, Wu Y, Jia G, Anaya E, Zhang J, et al. A CRISPR-Based Screen Identifies Genes Essential for West-Nile-Virus-Induced Cell Death. Cell Rep. 2015;12(4):673–83. Epub 20150716. doi: 10.1016/j.celrep.2015.06.049. PubMed PMID: 26190106; PubMed Central PMCID: PMCPMC4559080.

26. Savidis G, McDougall WM, Meraner P, Perreira JM, Portmann JM, Trincucci G, et al. Identification of Zika Virus and Dengue Virus Dependency Factors using Functional Genomics. Cell Rep. 2016;16(1):232–46. Epub 20160621. doi: 10.1016/j.celrep.2016.06.028. PubMed PMID: 27342126.

27. Song Y, Huang H, Hu Y, Zhang J, Li F, Yin X, et al. A genome-wide CRISPR/Cas9 gene knockout screen identifies immunoglobulin superfamily DCC subclass member 4 as a key host factor that promotes influenza virus endocytosis. PLoS Pathog. 2021;17(12):e1010141. Epub 2021/12/07. doi: 10.1371/journal.ppat.1010141. PubMed PMID: 34871331; PubMed Central PMCID: PMCPMC8675923.

28. Heaton BE, Kennedy EM, Dumm RE, Harding AT, Sacco MT, Sachs D, et al. A CRISPR Activation Screen Identifies a Pan-avian Influenza Virus Inhibitory Host Factor. Cell Rep. 2017;20(7):1503–12. Epub 2017/08/17. doi: 10.1016/j.celrep.2017.07.060. PubMed PMID: 28813663; PubMed Central PMCID: PMCPMC5568676.

29. Li Y, Muffat J, Omer Javed A, Keys HR, Lungjangwa T, Bosch I, et al. Genome-wide CRISPR screen for Zika virus resistance in human neural cells. Proc Natl Acad Sci U S A. 2019;116(19):9527–32. Epub 20190424. doi: 10.1073/pnas.1900867116. PubMed PMID: 31019072; PubMed Central PMCID: PMCPMC6510995.

30. Puschnik AS, Majzoub K, Ooi YS, Carette JE. A CRISPR toolbox to study virus-host interactions. Nat Rev Microbiol. 2017;15(6):351–64. Epub 2017/04/20. doi: 10.1038/nrmicro.2017.29. PubMed PMID: 28420884; PubMed Central PMCID: PMCPMC5800792.

31. Zhao C, Liu H, Xiao T, Wang Z, Nie X, Li X, et al. CRISPR screening of porcine sgRNA library identifies host factors associated with Japanese encephalitis virus replication. Nat Commun. 2020;11(1):5178. Epub 2020/10/16. doi: 10.1038/s41467-020-18936-1. PubMed PMID: 33057066; PubMed Central PMCID: PMCPMC7560704.

32. Hao J, Gao X, Light C, Sun Y, Lu S, Tian Y, et al. Genome-wide CRISPR/Cas9 knockout screen identifies host factors essential for Bovine Parainfluenza Virus Type 3 replication. bioRxiv. 2025.

33. Jin Z, Yi C, Zhou D, Wang X, Xie M, Zhou H, et al. Chicken genome-wide CRISPR library screen identifies potential candidates associated with Avian influenza virus infection. Int J Biol Macromol. 2025;293:139267. Epub 20241227. doi: 10.1016/j.ijbiomac.2024.139267. PubMed PMID: 39733882.

34. Han J, Perez JT, Chen C, Li Y, Benitez A, Kandasamy M, et al. Genome-wide CRISPR/Cas9 Screen Identifies Host Factors Essential for Influenza Virus Replication. Cell Rep. 2018;23(2):596–607. Epub 2018/04/12. doi: 10.1016/j.celrep.2018.03.045. PubMed PMID: 29642015; PubMed Central PMCID: PMCPMC5939577.

35. Schrader M, Reuber BE, Morrell JC, Jimenez-Sanchez G, Obie C, Stroh TA, et al. Expression of PEX11beta mediates peroxisome proliferation in the absence of extracellular stimuli. J Biol Chem. 1998;273(45):29607–14. doi: 10.1074/jbc.273.45.29607. PubMed PMID: 9792670.

36. Koch J, Pranjic K, Huber A, Ellinger A, Hartig A, Kragler F, et al. PEX11 family members are membrane elongation factors that coordinate peroxisome proliferation and maintenance. J Cell Sci. 2010;123(Pt 19):3389–400. Epub 20100907. doi: 10.1242/jcs.064907. PubMed PMID: 20826455.

37. Schrader M, Bonekamp NA, Islinger M. Fission and proliferation of peroxisomes. Biochim Biophys Acta. 2012;1822(9):1343–57. Epub 20111231. doi: 10.1016/j.bbadis.2011.12.014. PubMed PMID: 22240198.

38. Mann J, Reznik E, Santer M, Fongheiser MA, Smith N, Hirschhorn T, et al. Ferroptosis inhibition by oleic acid mitigates iron-overload-induced injury. Cell Chem Biol. 2024;31(2):249–64 e7. Epub 20231108. doi: 10.1016/j.chembiol.2023.10.012. PubMed PMID: 37944523; PubMed Central PMCID: PMCPMC10922137.

39. Cai Y, Liu H, Song E, Wang L, Xu J, He Y, et al. Deficiency of telomere-associated repressor activator protein 1 precipitates cardiac aging in mice via p53/PPARalpha signaling. Theranostics. 2021;11(10):4710–27. Epub 20210304. doi: 10.7150/thno.51739. PubMed PMID: 33754023; PubMed Central PMCID: PMCPMC7978321.

40. Robertson GL, Bodnya C, Gama V. Mitochondrial and peroxisomal fission in cortical neurogenesis. Int J Biochem Cell Biol. 2025;182–183:106774. Epub 20250328. doi: 10.1016/j.biocel.2025.106774. PubMed PMID: 40158688; PubMed Central PMCID: PMCPMC12136400.

41. He A, Dean JM, Lodhi IJ. Peroxisomes as cellular adaptors to metabolic and environmental stress. Trends Cell Biol. 2021;31(8):656–70. Epub 20210302. doi: 10.1016/j.tcb.2021.02.005. PubMed PMID: 33674166; PubMed Central PMCID: PMCPMC8566112.

42. Lodhi IJ, Semenkovich CF. Peroxisomes: a nexus for lipid metabolism and cellular signaling. Cell Metab. 2014;19(3):380–92. Epub 20140206. doi: 10.1016/j.cmet.2014.01.002. PubMed PMID: 24508507; PubMed Central PMCID: PMCPMC3951609.

43. Rizopoulos Z, Balistreri G, Kilcher S, Martin CK, Syedbasha M, Helenius A, et al. Vaccinia Virus Infection Requires Maturation of Macropinosomes. Traffic. 2015;16(8):814–31. Epub 20150506. doi: 10.1111/tra.12290. PubMed PMID: 25869659; PubMed Central PMCID: PMCPMC4973667.

44. Chu VC, McElroy LJ, Chu V, Bauman BE, Whittaker GR. The avian coronavirus infectious bronchitis virus undergoes direct low-pH-dependent fusion activation during entry into host cells. J Virol. 2006;80(7):3180–8. doi: 10.1128/JVI.80.7.3180-3188.2006. PubMed PMID: 16537586; PubMed Central PMCID: PMCPMC1440383.

45. Townsley AC, Weisberg AS, Wagenaar TR, Moss B. Vaccinia virus entry into cells via a low-pH-dependent endosomal pathway. J Virol. 2006;80(18):8899–908. doi: 10.1128/JVI.01053-06. PubMed PMID: 16940502; PubMed Central PMCID: PMCPMC1563910.

46. Bock C, Datlinger P, Chardon F, Coelho MA, Dong MB, Lawson KA, et al. High-content CRISPR screening. Nat Rev Methods Primers. 2022;2(1). Epub 20220210. doi: 10.1038/s43586-022-00098-7. PubMed PMID: 37214176; PubMed Central PMCID: PMCPMC10200264.

47. Michels BE, Mosa MH, Streibl BI, Zhan T, Menche C, Abou-El-Ardat K, et al. Pooled In Vitro and In Vivo CRISPR-Cas9 Screening Identifies Tumor Suppressors in Human Colon Organoids. Cell Stem Cell. 2020;26(5):782–92 e7. Epub 20200428. doi: 10.1016/j.stem.2020.04.003. PubMed PMID: 32348727.

48. Miles LA, Garippa RJ, Poirier JT. Design, execution, and analysis of pooled in vitro CRISPR/Cas9 screens. FEBS J. 2016;283(17):3170–80. Epub 20160616. doi: 10.1111/febs.13770. PubMed PMID: 27250066.

49. Schrader TA, Carmichael RE, Islinger M, Costello JL, Hacker C, Bonekamp NA, et al. PEX11beta and FIS1 cooperate in peroxisome division independently of mitochondrial fission factor. J Cell Sci. 2022;135(13). Epub 20220708. doi: 10.1242/jcs.259924. PubMed PMID: 35678336; PubMed Central PMCID: PMCPMC9377713.

50. Ferreira AR, Marques M, Ramos B, Kagan JC, Ribeiro D. Emerging roles of peroxisomes in viral infections. Trends Cell Biol. 2022;32(2):124–39. Epub 20211022. doi: 10.1016/j.tcb.2021.09.010. PubMed PMID: 34696946.

51. Villares M, Espert L, Daussy CF. Peroxisomes are underappreciated organelles hijacked by viruses. Trends Cell Biol. 2024. Epub 20241211. doi: 10.1016/j.tcb.2024.11.006. PubMed PMID: 39667991.

52. Ayres JS. Host-encoded antivirulence defenses: host physiologies teach pathogens to play nice. Curr Opin Immunol. 2024;91:102472. Epub 20241009. doi: 10.1016/j.coi.2024.102472. PubMed PMID: 39383570.

53. Li Z, Agellon LB, Allen TM, Umeda M, Jewell L, Mason A, et al. The ratio of phosphatidylcholine to phosphatidylethanolamine influences membrane integrity and steatohepatitis. Cell Metab. 2006;3(5):321–31. doi: 10.1016/j.cmet.2006.03.007. PubMed PMID: 16679290.

54. Cook KC, Moreno JA, Jean Beltran PM, Cristea IM. Peroxisome Plasticity at the Virus-Host Interface. Trends Microbiol. 2019;27(11):906–14. Epub 20190719. doi: 10.1016/j.tim.2019.06.006. PubMed PMID: 31331665; PubMed Central PMCID: PMCPMC6857447.

55. Ferreira AR, Marques M, Ribeiro D. Peroxisomes and Innate Immunity: Antiviral Response and Beyond. Int J Mol Sci. 2019;20(15). Epub 20190803. doi: 10.3390/ijms20153795. PubMed PMID: 31382586; PubMed Central PMCID: PMCPMC6695817.

